# Subcellular partitioning of *Nhlh2* mRNA reveals how *SNORD116* loss contributes to Prader-Willi Syndrome

**DOI:** 10.64898/2026.01.20.700624

**Authors:** Shadi Ariyanfar, Matthew A. Kocher, Christopher K. Thompson, Stuart Tobet, Deborah J. Good

**Author notes:** Lead Contact, @DeborahJGood. Instituto de Neurociencias CSIC-UMH, Spain. Departments of Biology and Neuroscience, St. Mary’s College of Maryland, St. Mary’s City, MD 20608.

## Abstract

Deletion of the *SNORD116* non-coding RNA is associated with the development of Prader-Willi Syndrome (PWS). We and others have identified *NHLH2/Nhlh2* as a putative *SNORD116/Snord116* target and in this study report that Nhlh2 and *Snord116* are co-expressed in forebrain neurons. *Nhlh2* mRNA is found predominantly in the nucleus of *Snord116+* hypothalamic neurons but is evenly distributed between the nuclear and cytoplasmic compartments in *Snord116-* neurons. A hypothalamic cell line co-expressing *Nhlh2* and *Snord116* recapitulates this nuclear partitioning pattern for *Nhlh2* only when the putative *Snord116* binding site is present in the 3’ untranslated region of the *Nhlh2* mRNA. In a PWS mouse model (Snord116^del^), *Nhlh2* mRNA levels are unchanged in *Pomc* neurons but are reduced and primarily cytoplasmic in lateral hypothalamic neurons. These findings demonstrate that *Snord116* regulates mRNA levels via mRNA partitioning, fundamentally altering our understanding of gene expression networks in PWS.

**Highlights:**

- *Snord116* snoRNA and its mRNA target *Nhlh2* are co-expressed in forebrain neurons.
- *Nhlh2* mRNA is predominantly nuclear co-localized in hypothalamic *Snord116+* neurons.
- A putative Snord116 binding site is within the region required for cellular partitioning.
- *Nhlh2* partitioning is lost in Snord116^del^ mice, a model of Prader-Willi Syndrome.

## Introduction

Prader-Willi syndrome (PWS) was first introduced in 1956 by Prader and colleagues^1^. It is a neurodevelopmental disorder of genomic imprinting, resulting from the absence of expression for the paternally active genes in a critical chromosomal region between 15q11.2-q13. In this region, the maternal genes are silent, and faulty genomic imprinting, aberrant DNA methylation, or deletion render the paternal genes silenced. Between 60 – 75% of PWS cases occur because of a *de novo* paternal genetic mutation of 15q11-q13^2–4^. In some of these cases, microdeletions remove just a portion of the region containing all or part of the *SNURF-SNRPN* transcript. Within this region, seven patients with PWS have microdeletions encompassing two C/D box snoRNAs coding regions, the SNORD116 cluster and SNORD 109A and the Imprinted in Prader-Willi exons (*IPW*)^5^. Interestingly, deletion of the whole cluster of Snord116 recapitulates the same core PWS phenotype of hyperphagia and obesity, developmental delay, facial characteristics, and excess weight gain. Thus, deletion or loss of *SNORD116* expression is critical to the development of PWS phenotypes. However, while *SNORD116*/*Snord116* (mouse) expression has been mapped to the central nervous system^6^, details of its expression pattern throughout the nervous system are incomplete. In particular, the identify of neurons within the hypothalamus that could be responsible for the phenotypes of hyperphagia, and later onset obesity in PWS are not known, with the exception of selected neuron types (e.g. NPY neurons,^7^).

*SNORD116* snoRNAs are derived from a long non-coding RNA transcript, *SNHG14*. In humans there are 30 annotated transcripts, numbered *SNORD116-1, SNORD116-2,* etc., with the entire locus referred to as *SNORD116@*^8^. While the *SNORD116@* snoRNAs contain a C/D box domain that is consistent with other SNORD RNAs, the biological function of this group of non-coding RNAs as well as mRNA and other targets that may interact with the non-coding RNAs has only recently begun to be elucidated. One of these targets, *NHLH2*, a basic helix-loop-helix transcription factor highly expressed in the hypothalamus, was originally shown to be significantly reduced in neurons differentiated from induced pluripotent stem cells derived from patients with PWS^9^. In addition, the *Nhlh2* knockout mouse displays greater phenotype-specific overlap with human PWS than all the PWS knockout mouse models^9–11^. Similarly, humans with missense mutations in *NHLH2* are hypogonadal, with delayed puberty and later-onset obesity^12^.

Our laboratory demonstrated that a putative *Snord116* binding site existed in the 3’ untranslated region (UTR) of *NHLH2/Nhlh2* which, when mutated, or when *Snord116* expression is absent, results in reduced levels of *Nhlh2* mRNA due to mRNA instability^13^ in a hypothalamic cell line (N29/2 cells^14^). While some studies reported low *Nhlh2* levels in ad libitum-fed and food deprived/refed animals^9^, a subsequent study failed to replicate low *Nhlh2* or *Pcsk1* mRNA levels in whole or micro-dissected hypothalamus^15^. In the current study, we analyzed *Nhlh2* mRNA in whole hypothalamic tissues of male and female WT and Snord116^m+/p-^ mice (PWS model), and in a knockdown study using Neuro2A cells. To define co-expression patterns of these genes in individual neurons, we carried out an extensive *in situ* hybridization study using RNAScope technology^16^. *Snord116@* is expressed in many forebrain areas^6^, while *Nhlh2* expression is highest in several regions of the hypothalamus, including the arcuate nucleus (ARC), paraventricular nucleus (PVN) and dorsal medial nucleus (DMH)^17–19^. The current study focused on *Pomc* and *Trh* neurons where *Nhlh2* is co-localized^20^. Whole body or Pomc neuron-specific targeted deletion of *Nhlh2* in mice lead to PWS-like phenotypes of obesity and pubertal delay^21,22^. These findings further clarify the spatial and temporal contexts in which *Nhlh2* is co-expressed with *Snord116@* noncoding RNAs. Most importantly, these studies reveal a novel subcellular localization pattern for *Nhlh2* mRNA in the presence versus absence of *Snord116@* expression that highlights a new role for *Snord116@*, and *SNORD* RNAs in general, in subcellular regulation of its mRNA targets.

## Results

### In situ hybridization experimental design, and confirmation

The *Snhg14, Snord116@*, and *Nhlh2* probe sets were designed by RNAScope™ (ACD Biotechne, Newark, CA) and the range of the probe sets for each are shown (**Figure 1A-C**). Note that the *Snord116* probe set that we used was designed to overlap the Snord116 mouse snoRNAs, which are similar in sequence for each snoRNA over the entire region of the *Snord116* locus^8^. Clusters 1 and 2 of the murine *Snord116@* locus contain 71 Snord116 transcripts, with approximately 84-88% homology to the group 1 human SNORD116 ncRNAs. In comparison, humans have only 30 annotated SNORD116 transcripts^8^. **Figures 1D-1E** show a representative area from N29/2 cells transfected with the *Nhlh2-myc* plasmid. Note the overlap between the *Nhlh2* probe and the myc probe in transfected cells compared to the lack of overlap, as well as relatively faint, *Nhlh2* signal in the non-transfected cells. Finally, we wanted to confirm that there was overlap between the *Snhg14* and *Snord116@* probes, and that we could detect only *Snhg14*, and not *Snord116@* in Snord116^del^ mice that have a homozygous deletion in both loci for the *Snord116* locus. Neurons from a medial coronal section of brain are shown for an aged matched male WT (**Figure 1F**) and *Snord116^del^* animal (**Figure 1G**). The *Snord116^del^* animal has both alleles of *Snord116@* deleted. The *Nhlh2* signal (red), the *Snhg14* (white), and *Snord116@* (green) signals were co-localized in the same cells, surrounding the nuclei (blue) for the WT mice, but only the *Snhg14* (white) signal with weak expression of *Nhlh2* could be detected in the *Snord116^del^* mouse. For the WT mice, we found a positive correlation (*P*< 0.001; 375 neurons) between *Snord116@* and *Snhg14* expression in the lateral hypothalamus.

**Figure 1:**
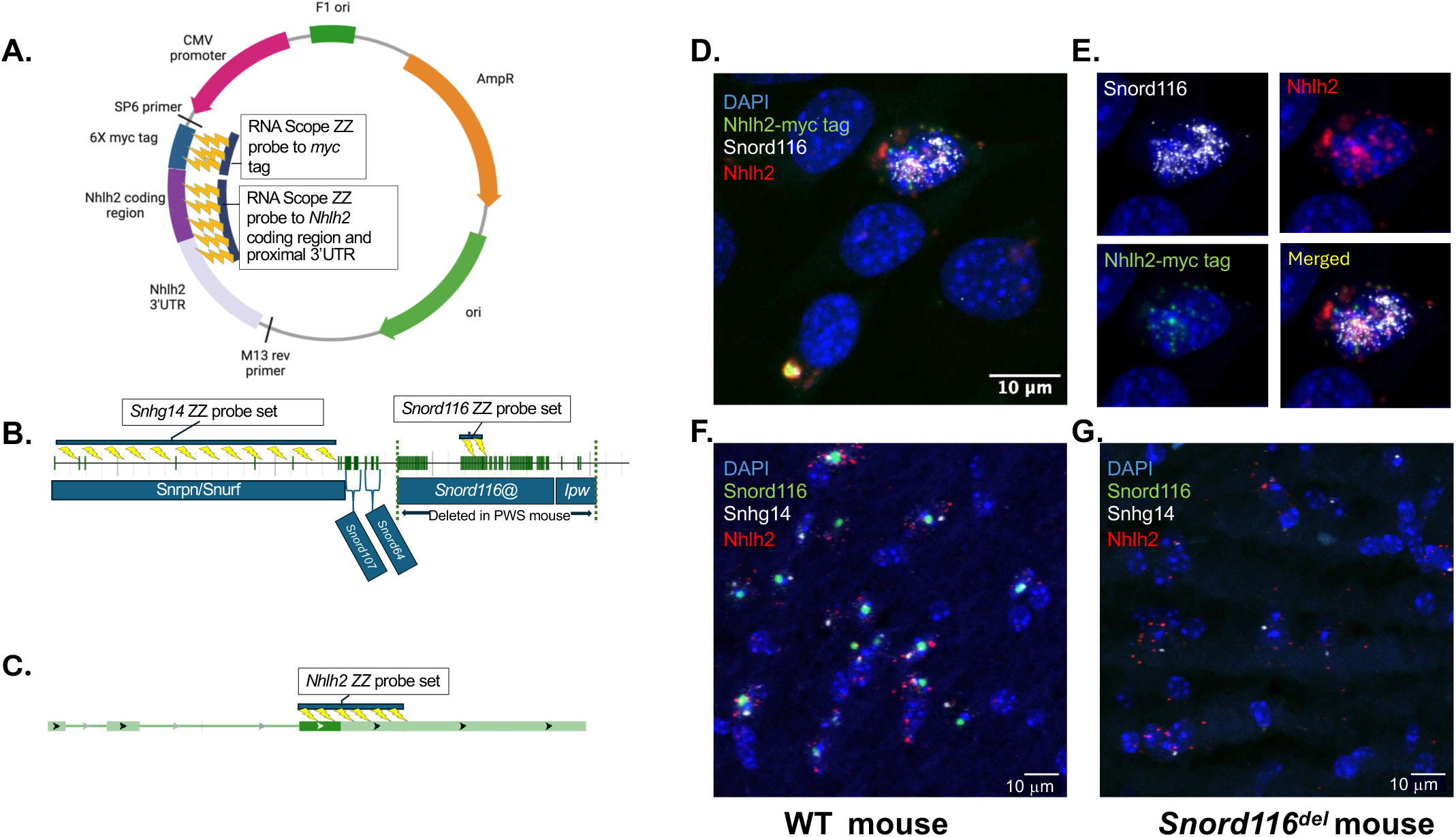
*Nhlh2, Snhg14,* and *Snord116* RNAScope Probes and Target areas. All the probes in this study were designed by Advanced Cell Diagnostics, Newark, CA. **A.** Graphical illustration of *Nhlh2-myc* plasmid and RNAScope probes to the myc tag region for specificity to transfected *Nhlh2*. The coding region probe will recognize both endogenous and transfected *Nhlh2.* **B.** The mouse Snhg14 host gene on mouse chromosome 7B5 and associated loci. The intron/exon structure of the mouse gene was obtained from MGI, ID 1289201. *Snhg14* and *Snord116* probe sets are marked on the graphical model. *Snord116* model probes focus mainly on group 1 *Snord116*. **C.** Graphical illustration of *Nhlh2* mRNA probe in the 3‘tail region. According to the probe set location, the hybrid mRNA gene can still be detected in the N2KO model. **D.** Validation of myc tag probe in transfected N29/2 cells using in situ hybridization RNAScope assay. Nhlh2 mRNA (CY5.5 red), myc tag (GFP, green), and Snord116 (TRITC, gray) probe shown. **E.** Channel-based visualization of individual probes. Nuclei counterstained with DAPI (blue). **F.** Validation of *Snhg14, Snord116*, and *Nhlh2* probe in tissues using in situ hybridization RNAScope assay. Representative 63x magnification image of a coronal forebrain section from WT (**F**.) and Snord116KO (**G.**) male mice. Nhlh2 mRNA (CY5.5 red), Snhg14 (TRITC, gray), and Snord116 (GFP, green) probe shown. Three representative cells are shown and acquired using a Leica SP8 confocal microscope. Nuclei counterstained with DAPI (blue).

### The *Nhlh2* mRNA 3’UTR controls nuclear partitioning in the presence of *Snord116* expression

We previously showed that *Nhlh2* mRNA stability was improved in the presence of *Snord116* expression using the hypothalamic cell line N29/2^13^. In the presence of a *Snord116* expression vector, Nhlh2 protein levels are increased only when an *Nhlh2* expression construct contains its full-length 3’ UTR, which includes a predicted *Snord116* binding site^13^. Like previously published studies, N29/2 cells were transfected with an *Nhlh2* myc-tagged expression vector containing either the full length 3’UTR or a partial 3’UTR missing the predicted *Snord116* binding site in the presence or absence of the *Snord116* expression vector. The myc-*tagged* Nhlh2 vector was used to accurately identify exogenous versus endogenous *Nhlh2* mRNA in this study. Since N29/2 cells do not express high levels of endogenous *Snord116@* the cell line served as a good model for PWS-like cells. RNAScope provides intracellular localization of *Nhlh2* and *Snord116* RNAs. As shown in **Figure 2, A-D**, in the presence of the *Snord116-3* vector, the *Nhlh2* mRNA containing the full length 3’UTR is localized primarily to the nuclear compartment, compared to the construct expressing the partial or short 3’UTR, where *Nhlh2-myc* is found in the cytoplasm and has a low expression level overall. Statistical analysis of individual puncta (45-120 neurons, expressing *Nhlh2-myc*, **Supplemental Tables 1-2**) shows that there are significantly fewer total *Nhlh2*-myc puncta for the full-length tail construct in the absence of *Snord116-3* expression, similar to what we previously observed using QPCR analysis^13^ (**Figure 2E, left panel**). This difference is pronounced when the amount of nuclear-localized *Nhlh2* message with the full-length 3’UTR is compared between the *Snord116-3* transfected versus empty vector transfected lines (**Figure 2E, middle panel**), while there is no significant difference in the total cytoplasmic levels of *Nhlh2* mRNA (**Figure 2E, right panel**). For the partial 3’UTR construct, *Nhlh2* expression levels are similar in total abundance and subcellular localization, regardless of *Snord116-3* expression (**Figure 2F**).

**Figure 2:**
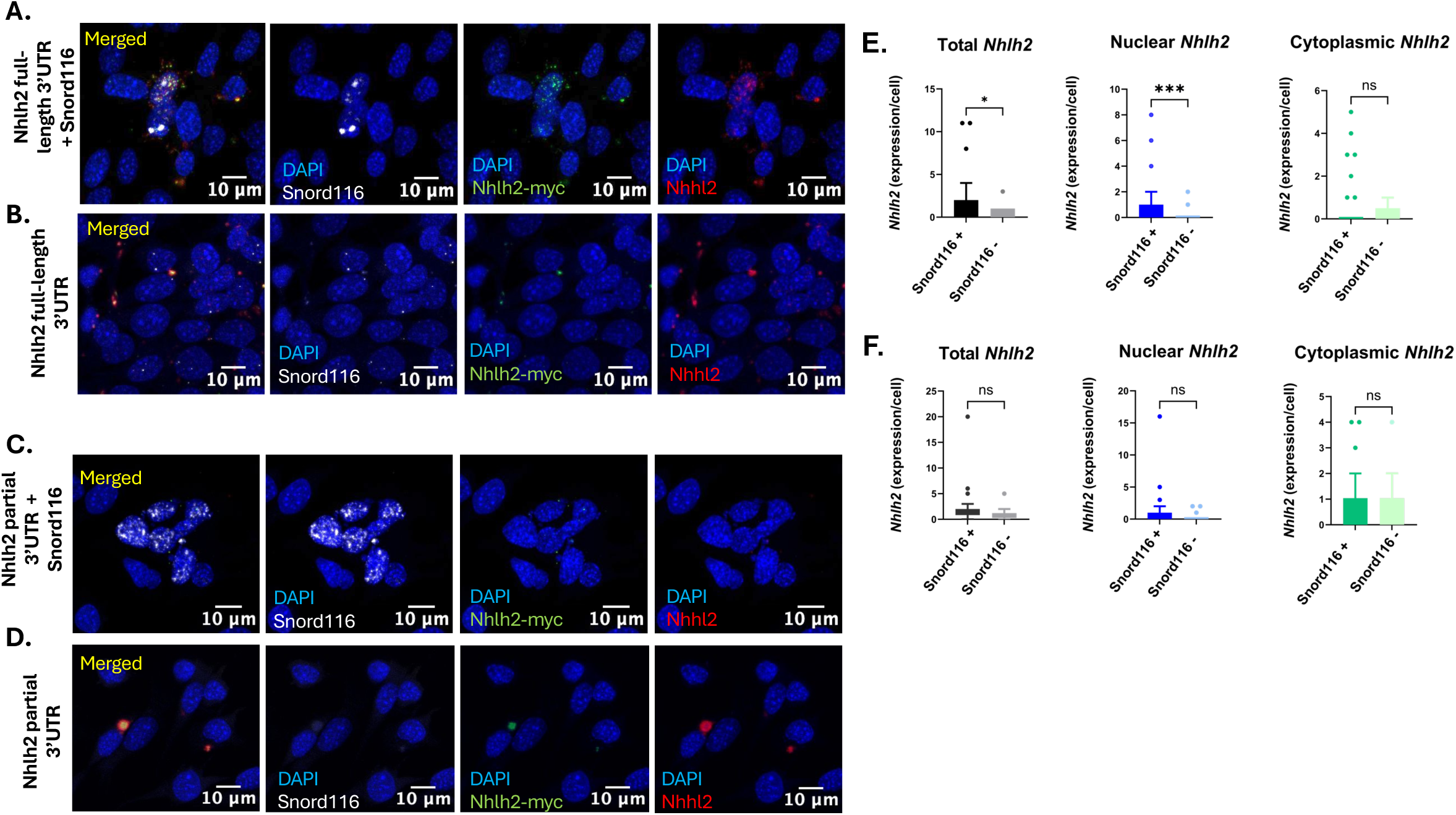
Analysis of *Nhlh2* expression in the mouse hypothalamus neuron cell line, N29/2. **A-B.** Representative 60x images of N29/2 hypothalamic neurons, transfected with a full-length *Nhlh2* 3’-UTR expression vector, containing a myc-tagged Nhlh2 (mouse) coding region and with (**A.**), or without (**B.**) a mouse Snord116 consensus expression plasmid. Cells were probed for *Nhlh2* myc-tagged motif (green), endogenous and exogenous *Nhlh2* (red), and *Snord116* (gray) using RNAscope in situ hybridization assay. Nuclei were counterstained with DAPI (blue). Panel is represented in a channel-based view from left to right**. C-D.** Representative 60x images of N29/2 hypothalamic neurons, transfected with a short-length *Nhlh2* 3’-UTR expression vector + *Snord116* (**C**.) or without *Snord116* (**D.**) Nuclei were counterstained with DAPI (blue). Panel is represented in a channel-based view from left to right. **E-F**. Quantification of Total, Nuclear, and Cytoplasmic expression levels for the full-length Nhlh2 3’-UTR expression vector and the partial (short) Nhlh2 3’ UTR expression vector. All graphs indicate median and IQR, and individual dots represent single cells with Nhlh2 counts. Whiskers extend to 1.5× IQR. Group differences were assessed using a two-tailed Mann–Whitney U test for transfection models.

### QPCR analysis of *Nhlh2, Psck1/3*, and *Snord116@* expression in WT versus *Snord116^m+/p^ mice*

The *Snord116^m+/p-^* mouse line^11^ has been used as a model for the minimal deletion of *Snord116* locus in some PWS patients. Given previous studies showing reduced overall expression of *Nhlh2* and *Pcsk1/3* levels in male Snord116^m+/p-^ animals, we sought to analyze *Nhlh2* and *Pcsk1/3* expression in this line in both sexes. As shown in **Figure 3**, we were unable to detect differences in *Nhlh2* or *Pcsk1/3* expression using this method for either males (**Figure 3A**) or for estrous-cycle matched females (**Figure 3B**).

**Figure 3:**
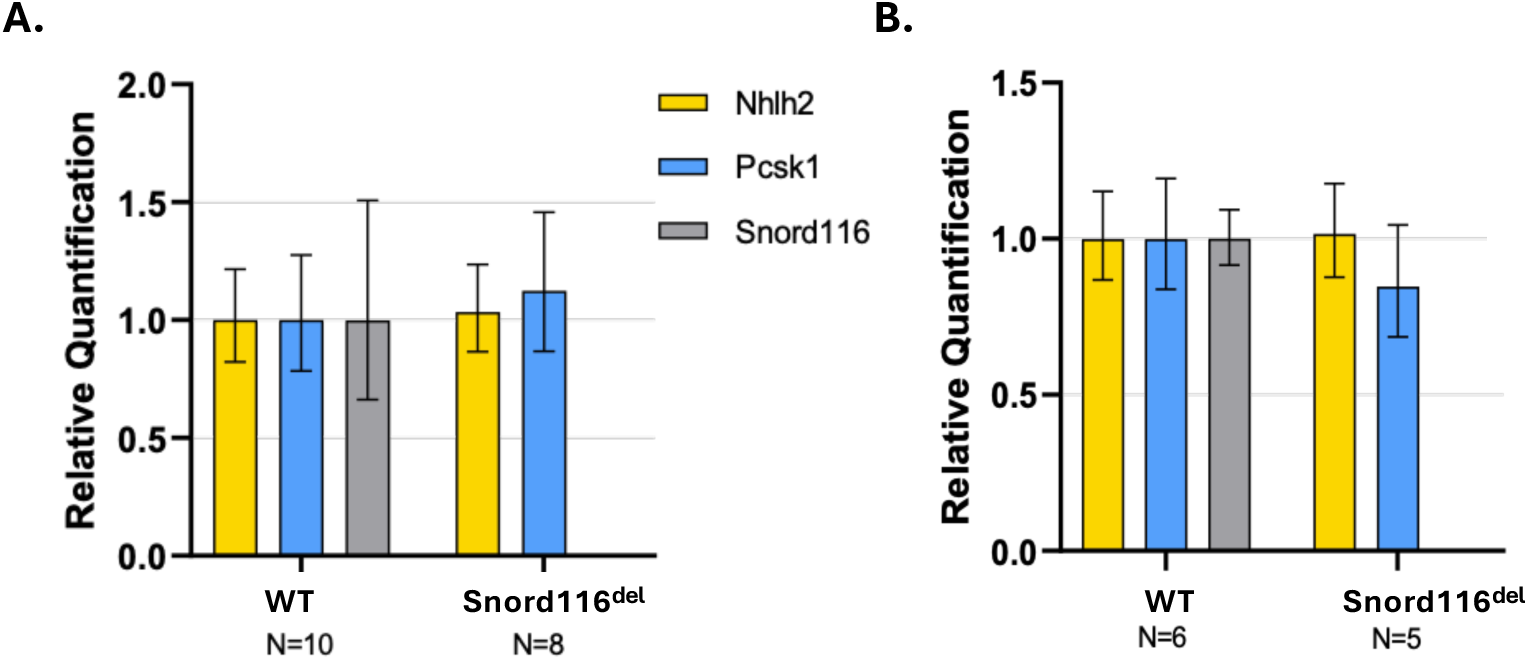
Whole hypothalamic expression of *Nhlh2, Pcsk1/3* and *Snord116*. RT-QPCR analysis of RNA isolated from whole hypothalamic blocks obtained from males (**A.**) or females (**B.**) Male and female WT mice, and mice with a paternal deletion of *Snord116* (Snord116^del^) were ad lib fed and were euthanized between 1-2 PM. Animal numbers for each variable is shown. Data shown is mean +/- standard deviation.

### Co-localization of *Snhg14*/*Snord116@* RNA with *Nhlh2* mRNA across forebrain regions

Previous work has shown *Snord116* expression throughout the mouse and human brains, and separately, *Nhlh2* mRNA expression in mouse brain, but the co-localization between these two RNAs was not known^2,6,10,19,23^. The findings in the neuronal cell-based studies, compared to the QPCR analysis of brain homogenates, led us to hypothesize that specific neuron types or brain regions would show reduced *Nhlh2* mRNA in the absence of *Snord116*, even when whole hypothalamic QPCR analysis did not. Thus, major subregions of the brain and spinal cord were individually characterized visually and quantitatively for co-expression patterns.

*Snhg14/Snord116@* is expressed in multiple forebrain regions, along with co-localized *Nhlh2*. The top row, (**panels 4A, D, G, J, M**) images at lower power show the regions of the hippocampus, habenula, cortex, thalamus, and hypothalamus. Higher magnification in the middle row (**panels 4B, E, H, K, N**) demonstrates that levels of each RNA vary by brain region. Using confocal imaging, within individual neurons (**panels 4C, F, I, L, O**), one can see that *Snord116@* and *Nhlh2* levels are much lower in the cortex, in which each neuron has only between 2-3 puncta for each message. Conversely, in the habenula, *Nhlh2* levels are visually much higher in the medial habenula, compared to the lateral habenula and *Snhg14* levels appear similar throughout the habenular regions. Quantitative analysis found that there can be up to 15 puncta for *Nhlh2* message in neurons that have 0-4 puncta for *Snhg14*.

To statistically evaluate the relationship between *Snhg14*/*Snord116@* and *Nhlh2* expression across various brain regions, we performed negative binomial regression for analysis of all regions. In the dentate gyrus (**Supplemental Figure 1A,** p < 0.0001) a statistically significant positive association between *Snhg14* expression and *Nhlh2* expression per nucleus in the dentate gyrus was observed, with *Snhg14* expression significantly predicting *Nhlh2* expression corresponding to a 1.78-fold (78%) increase in *Nhlh2* counts per unit increase in *Snhg14* (**Supplemental Table 3-4**)). A similar association pattern is found for the medial habenula (**Supplemental Figure 1B**, p < 0.0001, **Supplemental Tables 5-6**), the cortex (**Supplemental Figure 1C**, p < 0.0001, **Supplemental Tables 7-8**), and the thalamus (**Supplemental Figure 1D,** p < 0.0001, **Supplemental Tables 9-10**).

In the paraventricular nucleus (PVN) and the arcuate nucleus (ARC) of the hypothalamus the model revealed a significant positive relationship between *Snhg14* and *Nhlh2* expression. There was a significant positive association between nuclear *Snhg14* expression and *Nhlh2* expression per nucleus in the PVN (**Figure 4O**, *P* < 0.0001, **Supplemental Table 11**) where *Snhg14* expression was a statistically significant positive predictor of *Nhlh2* expression per nucleus corresponding to a 12% increase in the expected Nhlh2 count with *Snhg14* expression (**Supplemental Table 12**). There was a significant positive association between nuclear *Snhg14* expression and *Nhlh2* expression per nucleus in the ARC (**Figure 4N**, **Supplemental Figure 1E**, *P* = 0.0002; **Supplemental Tables 13-14**), corresponding to a 59.1% increase in the expected count of *Nhlh2* puncta for each unit increase in *Snhg14* expression.

**Figure 4:**
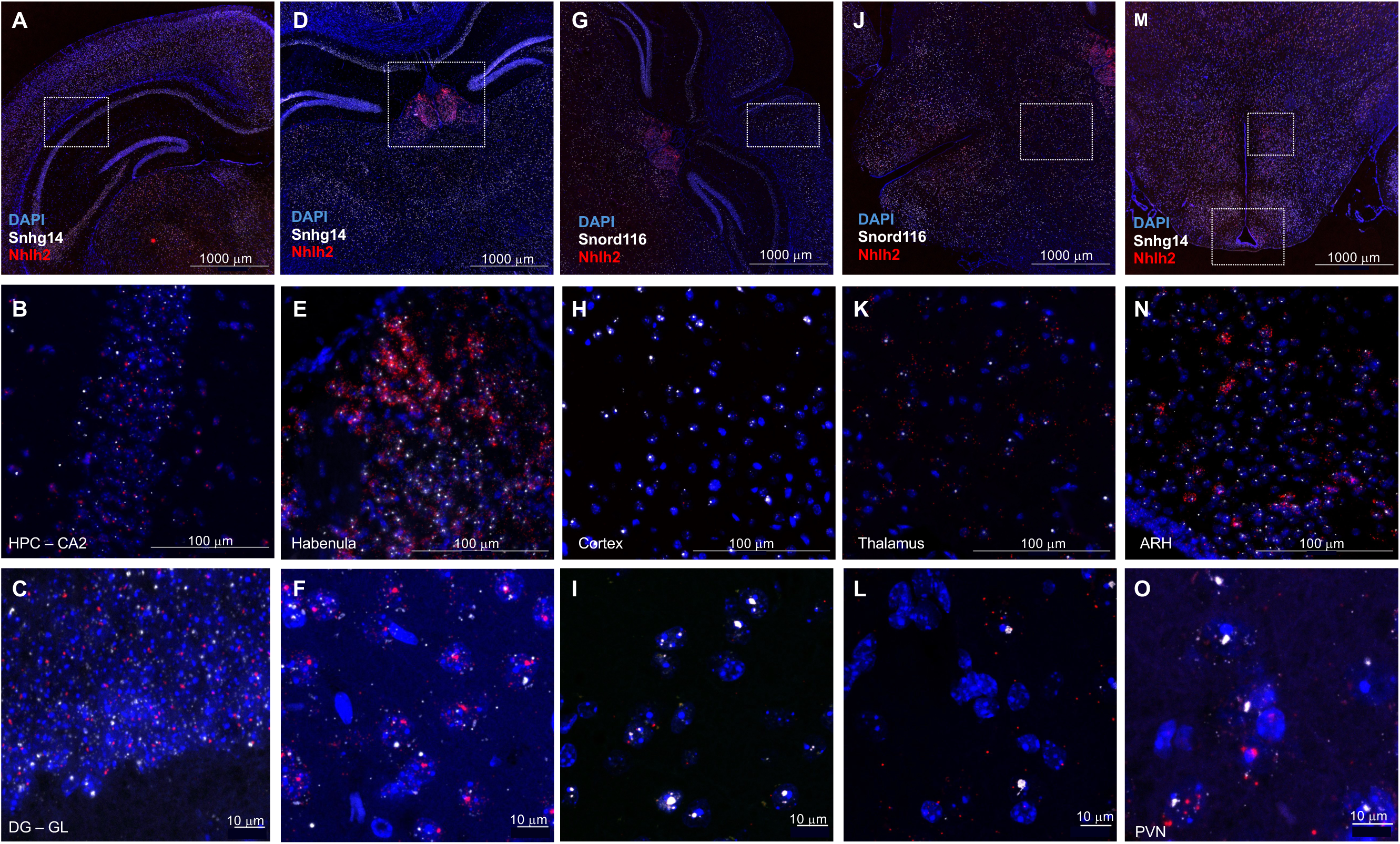
Region-specific colocalization of Snhg14/Snord116 and Nhlh2 mRNA in adult WT 129Sv/J forebrain. RNAscope in situ hybridization was used to identify cells co-expressing Nhlh2 and Snhg14/Snord116 in critical brain regions responsible for different phenotypes of PWS. Different regions are probing for Nhlh2 mRNA (red), and Snhg14/Snrod116 (white), acquired from Cytation 5 imaging reader, counterstained with DAPI (blue) in pseudo color. **A.** Representative 4x visualization of the hippocampus and dentate gyrus. **B**. Shows 60x visualization of the hippocampus Cornu Ammonis (CA)2. **C**. Representative 63x magnification of the granular layer of the dentate gyrus. **D.** Representative 4x visualization of the habenula. **E.** Shows 60x visualization of the medial habenula **F.** Representative 63x magnification of medial habenula. **G.** Representative 4x visualization of the cortex. **H**. Shows 60x visualization of cortex. **I.** Representative 63x magnification of the cortex **J.** Representative 4x visualization of the thalamus. **K.** Representative 60x visualization of the thalamus. **L.** Representative 63x visualization of the thalamus. **M.** Representative 4x visualization of the hypothalamic nuclei. **N.** Representative 60x magnification of the arcuate nucleus. **O.** Representative 63x visualization of paraventricular hypothalamic nucleus. All 4x and 60x magnified images acquired from Cytation 5 imaging multimode reader converted microscope. 63X magnification images acquired using Confocal microscopy using identical image acquisition settings (laser power and gain) across all sections in each experiment from each tissue type. The images were then colorized, z-projected, and prepared using identical contrast and brightness parameters in ImageJ.

The findings support a robust, nonlinear effect of *Snhg14* expression on *Nhlh2* expression at a single-cell resolution, demonstrating that *Snhg14/Snord116@* expression is a statistically significant and biologically meaningful predictor of *Nhlh2* expression in individual neurons. The nonlinear, multiplicative relationship observed here suggests a potential regulatory or co-expressive role for *Snhg14* in shaping *Nhlh2* mRNA levels at the single-cell level.

The study thus far demonstrates that *Nhlh2* mRNA co-localizes in neurons with *Snhg14/Snord116@* selectively in the nuclear compartment, rather than the cytoplasmic compartments, where many mRNAs are localized once spliced. While the RNAScope probe for *Nhlh2* would not allow us to differentiate between spliced and unspliced *Nhlh2* mRNA, the pattern observed was still different than what was expected for most mRNAs.

To characterize the subcellular localization pattern further, *Pomc* and *Trh* expressing neurons in the hypothalamus were co-localized with *Nhlh2* and *Snhg14/Snord116* probes. Confocal microscopy, either alone to demonstrate the individual z-steps through individual neurons, as well as in conjunction with the Imaris™3D image analysis software, was used to localize individual mRNAs and ncRNAs. *Pomc* mRNA (green) was found almost exclusively in the cytoplasmic compartment. As shown in **Figure 5A-F**, in both *Pomc*-expressing (green) and non-expressing neurons, *Nhlh2* signal (red) can be found predominantly in the nucleus, co-localized with *Snhg14* and the DAPI nuclear/nucleolar counterstain. *Pomc* mRNA (green) was found almost exclusively in the cytoplasmic compartment.

**Figure 5.**
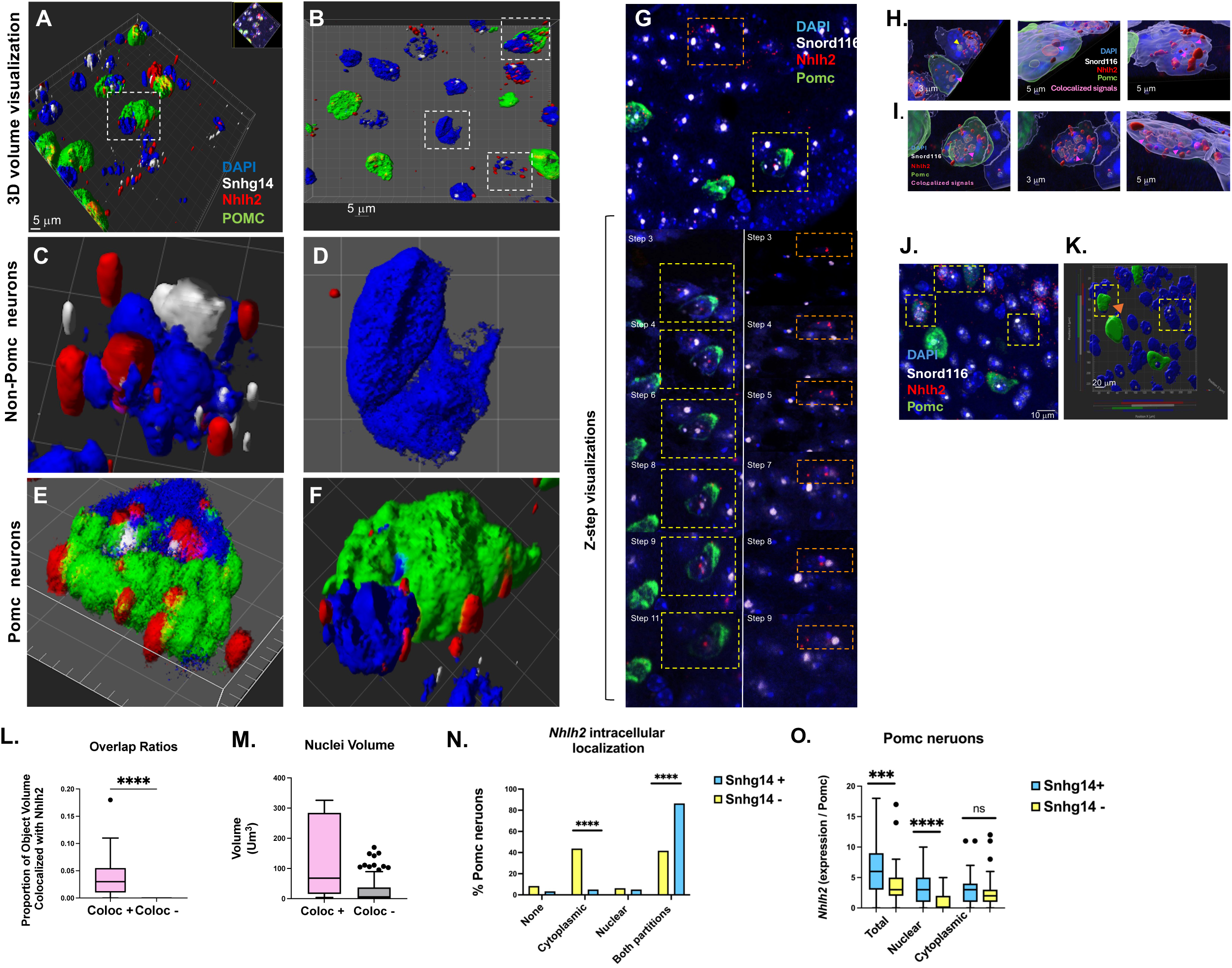
Single cell analysis of caudal arcuate Pomc neurons focusing on the Spatial engagement of Snhg14 and Nhlh2 pattern. **A.** Representative 3D image of RNAscope immunofluorescent staining of caudal arcuate Pomc neurons. **B.** Representative 3D image of a single Pomc and non-Pomc neuron. **C.** Representative 3D image of RNAscope immunofluorescent staining of caudal arcuate neuron with Snhg14 (gray) and Nhlh2 (red) nuclear expression. **D.** Representative 3D visualization of a single neuron of ARC with no detected Snhg14 expression. **E.** Representative 3D visualization of a single Pomc neuron of arcuate, pointing at nuclear co-expression of Snhg14 and Nhlh2 mRNA in the nucleus. **F.** Representative 3D visualization of a single Pomc neuron of arcuate with no Snhg14 expression, showing partitioning of Nhlh2 mRNA in the cytoplasmic region. **G**. Cell-specific confocal Z-stack series of Pomc neuron (yellow box, left stack) and a non-Pomc neuron (orange box, right stack) of the caudal arcuate at 63x magnification. Each panel represents a single optical section taken at intervals of [0.75] µm along the Z-axis. **H-K**. Imaris-based 3D volume render colocalization analysis of Pomc neuron of the caudal arcuate nucleus, representing single-cell tracking analysis for Pomc neurons of Arcuate probing for *Snord116* (gray), *Nhlh2* (red), and Pomc (green), using manual thresholding. Nuclei stained with DAPI (blue). **H**. Shows Pomc and non-Pomc neurons with colocalization of *Snord116* and *Nhlh2* in the WT mouse model (left image). The colocalization signal (magenta) can be seen in the nucleus of both Pomc (middle image) and non-Pomc neurons (right image) with expression of *Snord116*. The overlapped signal in the nucleus is indicated by magenta arrowhead. **I**. Is a representative single Pomc neuron with colocalization of *Snord116* and *Nhlh2,* showing 3D view of Pomc with all 4 channels (left). The middle panel is a representative image of the same Pomc neuron from below the surface, where only the nucleus is visible, and the panel on the right is a representative image of the same Pomc without the green channel to confirm nuclear signals. **J**. Representative 63x magnification of the ARH from confocal microscopy used for 3D rendering. **K.** 3D colocalization analysis showing the overlapped signals on y and x axes indicating Snord116 (gray), Nhlh2 (red), Pomc (green), and DAPI (blue). **L**. Box-and-whisker plot showing quantitative colocalization analysis from Imaris, showing the overlap volume ratio between two fluorescent channels (Snord116 and Nhlh2). **M.** Box-and-whisker plot showing differences in volume of nuclei between colocalized and non-colocalized signals of interest. **N.** Stacked bar plots show differential distribution of localization across Pomc neurons of the caudal arcuate based on *Snhg14* expression. Fisher’s exact test revealed a highly significant difference in distribution across groups (P < 0.0001), indicating that subcellular localization is strongly associated with group identity. **O.** Box-and-whisker plots show total, nuclear, and cytoplasmic Nhlh2 signal counts in individual Pomc neurons grouped by Snhg14 expression status. Snhg14⁺ neurons (blue) exhibited significantly higher total (*P* = 0.0016) and nuclear (*P* = 0.0001) Nhlh2 expression compared to Snhg14⁻ neurons (yellow), as determined by Mann–Whitney U tests. Post hoc correction for multiple comparisons across subcellular compartments was applied using the Tukey-style method.

In non-*Pomc* neurons, *Nhlh2* was highly expressed and partitioned to the nucleus with *Snhg14* ncRNA (**Figure 5C**). However, *Nhlh2* mRNA was extremely low in those cells that did not express *Snhg14,* and in those cells, *Nhlh2* mRNA was more likely to be found in the cytoplasm (**Figure 5D**). The localization pattern is even more striking in *Pomc* neurons (green, **Figures 5E, F**), where *Nhlh2* mRNA was found in the nuclear compartment with *Snhg14* ncRNA, as well as in the cytoplasmic compartment with *Pomc* mRNA when *Snhg14* message was co-expressed (**Figure 5E**). *Nhlh2* mRNA puncta surrounded the nucleus or were found in the cytoplasm, colocalized with *Pomc* mRNA in the absence of *Snhg14* co-expression (**Figure 5F**).

To confirm that the observed pattern was to specific to individual neurons and not due to overlapping neuron cell bodies within the section, we performed z-step analysis on multiple examples where *Nhlh2* mRNA was found in the nuclear compartment. **Figure 5G** shows representative z-steps for one Pomc and one non-Pomc neuron in the same focal plane. There were steps where *Nhlh2* and *Snhg14* messages were found in the same 0.75 mM optical section (i.e. step 3, and 8 of the Pomc neuron; step 4 and 8 of the non-Pomc neuron), even with the signals overlapping one another.

### Nuclear and cytoplasmic localization of *Nhlh2* in hypothalamic neurons

For a more robust and accurate single-cell tracking analysis, a spatial colocalization assessment was performed. Using 3D volume rendering tool with manual thresholding detection, we were able to specifically detect overlapping signals between *Snord116@* and *Nhlh2* in Pomc and non-Pomc neurons. **Figure 5H** shows Pomc and non-Pomc neurons with colocalization of *Snord116* and *Nhlh2* in the WT mouse model. *Snord116@* ncRNA was detected as a pink hue where there was spatial colocalization and signal overlap with *Nhlh2* mRNA (indicated by the pink arrowhead and shown as a colocalized area in purple). The yellow arrowhead points at a non-colocalized *Snord116* signal when there was no merge between the two signals. (**Figure 5H**, left image). A single Pomc neuron with overlapping *Snord116@* and *Nhlh2* signal is shown in **Figure 5H**, middle panel, while the right panel shows the adjacent non-POMC neuron with colocalization detected (pink arrowheads). The results in **Figure 5I** confirm nuclear colocalization of *Snord116@* and *Nhlh2* in a Pomc neuron with all 4 channels (left and middle). The middle image is the same cell viewed from the bottom surface, where only the nucleus is visible, and the image on the right shows the same view with the green channel (Pomc) turned off to confirm the nuclear colocalization signals for both RNAs. **Figure 5J** shows a confocal image of the representative neurons (yellow boxes) used to visually confirm nuclear co-localization of *Nhlh2* mRNA and *Snord116@* snoRNA in **Figure 5I and H**. Most of the DNA regions in this stacked confocal image contain *Snord116* (grey). **Figure 5K** shows a three-dimensional surface rendering model of the area in **Figure 5H-J**, generated by the Imaris™ software, and shows a projection bar (X-axis, bottom) with colored segments representing the signal overlaps. The majority of the *Nhlh2* mRNA puncta (red) were found within and around the DNA regions (blue), and overlap with *Snord116* puncta (grey, X-axis). This difference was strongly supported by a Mann–Whitney U test (U = 803.0, p < 2 × 10⁻¹⁶) and confirmed by Welch’s t-test (t = 2.82, *P* = 0.017), indicating that overlap is a defining feature of the colocalized population **(Figure 5L, Supplemental Table 15)**. Analysis of object volume and colocalization revealed consistent differences between *Snord116@* and *Nhlh2* colocalized (Coloc⁺) and non-colocalized (Coloc⁻) groups. Absolute volume measurements showed that Coloc⁺ nucleus (DAPI) were significantly larger (mean = 122.66) compared to Coloc⁻ objects (mean = 29.72; t-test, p = 0.03), with substantially greater variance confirmed by F-test (p < 0.01). Distributional differences were further supported by the Kolmogorov–Smirnov test (p = 0.03) **(Figure 5M, Supplemental Table 16).**

We examined subcellular localization patterns of *Nhlh2* in two groups of Pomc neurons stratified by Snhg14 expression status: Snhg14⁺ (n = 41 cells) and Snhg14⁻ (n = 35 cells). Cells were classified into four *Nhlh2* mRNA localization categories: None, Cytoplasmic, Nuclear, and Both partitions (nuclear and cytoplasmic). Among all Pomc neurons analyzed, nuclear *Nhlh2* localization was relatively rare, occurring in 5% of Snhg14⁺ cells and 6% of Snhg14⁻ cells. In contrast, cytoplasmic localization was more frequent in the Snhg14⁻ group (44%) than in the Snhg14⁺ group (5%). When *Nhlh2* mRNA was detected in both nuclear and cytoplasmic compartments, this pattern was strongly enriched in the Snhg14⁺ group (87%) compared with the Snhg14⁻ group (42%), indicating that the localization of *Nhlh2* mRNA is preferentially associated with *Snhg14* expression as absence of *Nhlh2* mRNA or cytoplasmic restriction of *Nhlh2* mRNA is predominantly observed in *Snhg14*- cells **(Figure 5N, Supplemental Tables 17-18**, P < 0.001). Next, across all biological replicates, we evaluated individual group differences using Fisher’s exact tests of *Nhlh2* subcellular localization patterns in *Snhg14*+ and *Snhg14*− neurons. The frequency of cells lacking a detectable *Nhlh2* signal (None) did not differ between groups *(P* = 0.4048), indicating comparable baseline detection across conditions. In contrast, cytoplasmic *Nhlh2* localization was markedly enriched in *Snhg14*− neurons (*P* < 0.0001), reflecting a strong shift toward cytoplasmic accumulation in the absence of *Snhg14* expression. Nuclear localization remained low in both groups and did not differ significantly (P > 0.9999). Conversely, dual nuclear + cytoplasmic localization (“Both”) was significantly overrepresented in *Snhg14*+ neurons (*P* < 0.0001), suggesting that *Snhg14* expression is associated with higher nuclear access and/or retention of *Nhlh2* mRNA. Together, these analyses demonstrate a clear redistribution of *Nhlh2* localization depending on *Snhg14* status, with *Snhg14*+ neurons strongly favoring dual-compartment localization and *Snhg14*− neurons disproportionately accumulating *Nhlh2* in the cytoplasm. This suggests that the subcellular distribution of *Nhlh2* may be influenced by the expression of *Snhg14* (**Supplemental Tables 19- 22).**

Analysis of *Nhlh2* mRNA counts between *Snhg14*⁺ and *Snhg14*⁻ neurons within the Pomc subcellular compartments showed that total *Nhlh2* puncta counts were significantly higher in *Snhg14*⁺ neurons compared to *Snhg14*⁻ neurons (median 6 vs. 3; *P* = 0.0001), indicating an overall elevation of *Nhlh2* mRNA in the presence of *Snhg14*. This effect was driven primarily by nuclear Nhlh2 enrichment: nuclear puncta were markedly increased in *Snhg14*⁺ neurons (median 3 vs. 0; *P* < 0.0001), consistent with a model in which *Snhg14* facilitates nuclear localization or stability of *Nhlh2* transcripts. In contrast, cytoplasmic *Nhlh2* levels did not significantly differ between groups (median 3 vs. 2; *P* = 0.2830), suggesting that cytoplasmic localization or retention of *Nhlh2* is not substantially altered by *Snhg14* status (**Supplemental Table 23**). Together, these analyses reveal that *Snhg14* primarily influences the nuclear localization and overall abundance of *Nhlh2* rather than cytoplasmic accumulation alone.

*Nhlh2* and *Snhg14* visually colocalized in Trh neurons and non-Trh neurons in DMH (**Figure 6A, B)**. A Z-stack series confirms the overlapping signals **(Figure 6C)** with an analysis of 73 Trh neurons revealing a significant association between Snhg14 expression and *Nhlh2* localization patterns (*P* = 0.0361; **Supplemental Tables 24-25).** However, none of the individual localization categories reached statistical significance when tested independently (**Figure 6E**; *P* > 0.05). These data indicate that *Snhg14*+ Trh neurons tend to have a broader subcellular distribution of *Nhlh2,* showing that the distribution was not strongly dependent on Snhg14 expression in Trh neurons, with no single compartment showing a statistically robust shift between the two groups. **(Supplemental Tables 26-29).** Next, *Nhlh2* mRNA counts in Trh neurons were examined **(Figure 6F)**. Total *Nhlh2* counts were significantly higher in *Snhg14*+ neurons (median 6.5 vs. 3.5; *P* = 0.0299), indicating an overall increase in mRNA associated with *Snhg14* expression. Cytoplasmic *Nhlh2* levels were also elevated in *Snhg14*+ neurons (median 4 vs. 1; *P* = 0.0003). In contrast, nuclear *Nhlh2* counts did not differ significantly between groups (median 3 vs. 2; *P* = 0.6085), confirming that the nuclear accumulation of *Nhlh2* in Trh neurons was unaffected by *Snhg14* status. **(Supplemental Table 30)**.

**Figure 6:**
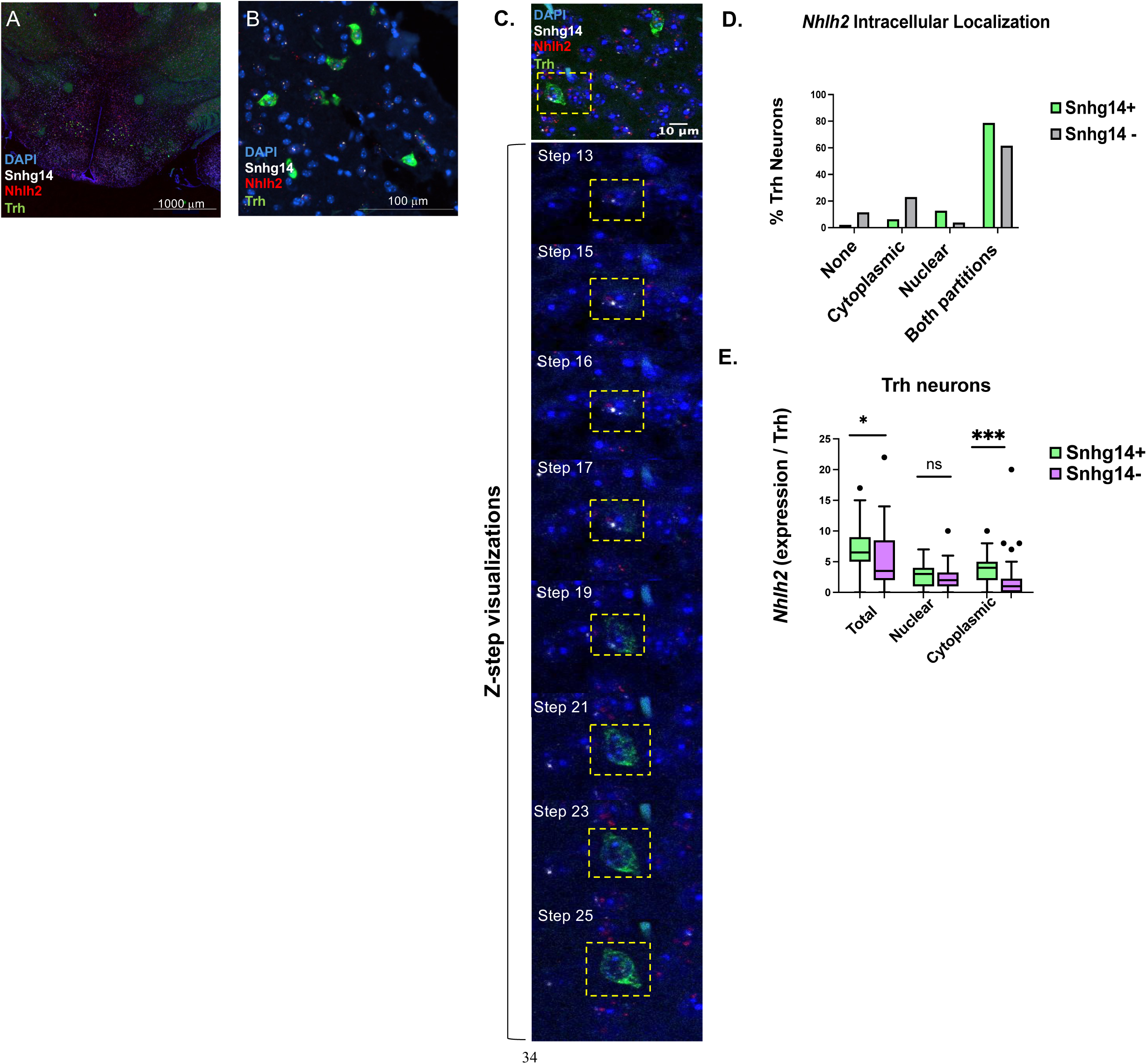
Single cell analysis of DMH Trh neurons. **A.** Regional representation of RNAscope in situ hybridization for *Snhg14* (white) and *Nhlh2* (red) in Trh neurons (green), counterstained with DAPI (blue). 4X magnification **B.** 60x magnification of Trh neurons, *Nhlh2*, and *Snhg14***. C.** Cell-specific confocal Z-stack series of Trh neurons of Dorsomedial hypothalamic nucleus at 63x magnification. Each panel represents a single optical section taken at intervals of [0.30] µm along the Z-axis. **E.** Stacked bar graph showing the distribution of Nhlh2 localization (None, Cytoplasmic, Nuclear, Both) in cells with (green) or without (gray) Snhg14 expression. Percentages represent the proportion of each localization type within Snhg14⁺ and Snhg14⁻ groups. Fisher’s exact test showed no significant association between *Nhlh2* localization and *Snhg14* status. **F.** Box-and-whisker plots show total, nuclear, and cytoplasmic *Nhlh2* signal counts in individual Trh neurons grouped by Snhg14⁺ (green) and Snhg14⁻ (magenta) cells across total, nuclear, and cytoplasmic compartments. Trh expression was significantly higher in the cytoplasm of Snhg14⁺ cells compared to Snhg14⁻ cells (*P* = 0.0003), but not at the nuclear and total count level as determined by two-tailed Mann–Whitney U tests. Bars represent medians and interquartile ranges; individual data points are shown. Post hoc correction for multiple comparisons across subcellular compartments was applied using the Tukey-style method.

### *Nhlh2* colocalization pattern differs between WT and Snord116^del^ animals

Mice with a deletion of both alleles of the *Snord116@* gene, termed Snord116^del^, were used to ensure that there was no leaky expression of *Snord116@* from the maternal allele. The lateral hypothalamus (LH), ARC, and DMH were characterized for *Snhg14/Snord116* and Nhlh2 co-expression patterns in WT and Snord116^del^ mice. As shown in **Figure 7A** (left panel), *Snord116* and *Snhg14* ncRNAs are co-expressed in WT LH with abundant *Nhlh2* mRNA. However, in Snord116^del^ mice (right panel), there were fewer puncta for *Nhlh2* mRNA, and no detection of the *Snord116* ncRNA. A single punctum for *Snhg14* message allows quantification between Snord116 expressing and non-expressing neurons. Levels of *Snhg14* expressing and non-expressing neurons remained identical between the two genotypes (**Figure 7B, Supplemental Table 32**). Quantification of *Nhlh2* signals indicates a significant reduction in *Nhlh2* individual puncta per cell in the Snord116^del^ mice versus WT animals only when *Snhg14* and *Nhlh2* were co-expressed (**Figure 7C, Supplemental Tables 33-34;** *P <0.001*). In the LH, this overall reduction translates to a significant reduction in total, nuclear, and cytoplasmic *Nhlh2* expression in the Snord116^del^ mice (**Figure 7 D-F, Supplemental Table 35,** *P* <0.0001).

**Figure 7:**
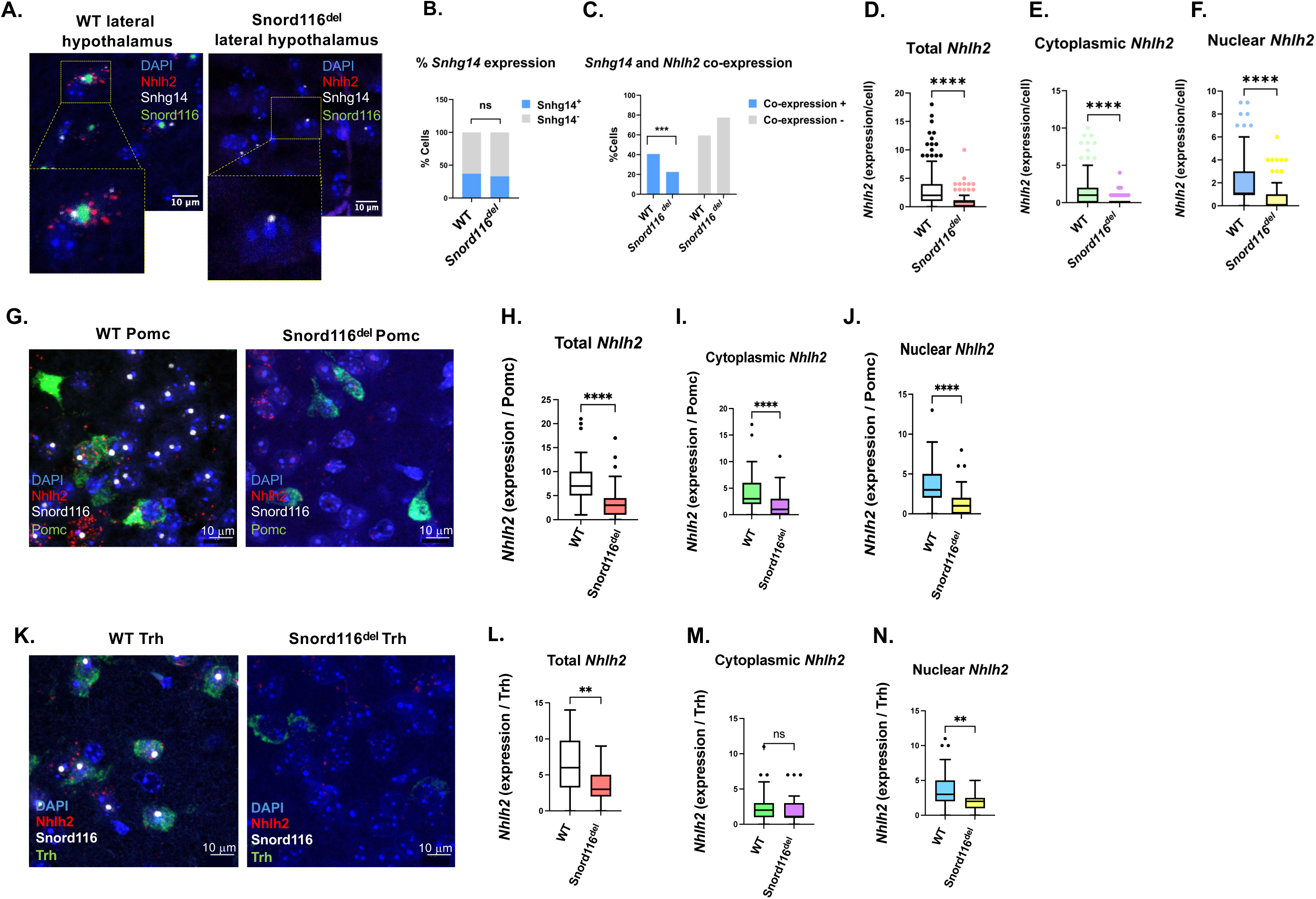
Subcellular co-expression of *Nhhl2* in WT and Snord116^del^ mouse model. **A**. Representative images of adult WT (left) and *Snord116^del^* (right) mice, showing lateral hypothalamic neurons probed for *Nhhl2* (red), *Snhg14* (green), and *Snord116* (gray) **B.** Quantification of total *Snhg14* as % counts. **C.** quantification of % neurons with *Snhg14* and *Nhlh2* co-expression in both models. Bar graph summarizing the % of cells expressing *Snhg14* in WT and *Snord116^del^* mice. Fisher’s exact test was used to show a significant association between genotype and *Snhg14* expression status. **D-F.** Quantification of total *Nhhl2* (**D**.), Cytoplasmic *Nhlh2* (**E**.), and Nuclear *Nhlh2* (**F**.) expression levels in LH neurons from both WT and Snord116^del^ mice. All graphs indicate median and interquartile range (IQR). **G.** Representative images of adult WT (left) and *Snord116^del^* (Right) mouse, showing Pomc-labeled neurons (green) probing for subcellular co-expression of *Nhhl2* (red) and *Snord116* (gray). **H-J.** Quantification of total *Nhlh2* (**H**.), Cytoplasmic (**I**.), and Nuclear (**J**.) expression level in Pomc neurons from both WT and Snord116^del^ mice. The bars indicate median and IQR. **K.** Representative images of adult WT (left) and *Snord116^del^* (Right) mouse, showing Trh-labeled neurons (green) probing for subcellular co-expression of *Nhhl2* (red) and *Snord116* (gray). **L-N.** Quantification of total *Nhlh2* (**L**.), Cytoplasmic (**M**.), and Nuclear (**N**.) expression level in Pomc neurons from both WT and Snord116^del^ mice. All graphs indicate median and IQR, and individual dots represent single cells with *Nhlh2* counts. Whiskers extend to 1.5× IQR. Group differences were assessed using a two-tailed Mann–Whitney U test for genotype. ∗*P* < 0.05, ∗∗*P* < 0.01, ∗∗∗∗*P* < 0.0001.

Representative Pomc neurons from the ARC region of the hypothalamus, both not expressing and expressing *Pomc* (green), appear to have visually fewer *Nhlh2* mRNA positive neurons (red) in the Snord116^del^ mice (**Figure 7G**). This trend can also be visualized with the video three-dimensional models (**Multimedia Video 1**). Quantification of these results shows that there is a significant decrease in total *Nhlh2* mRNA, as well as both cytoplasmic and nuclear localized *Nhlh2* mRNA in the Snord116^del^ mice compared to WT (**Figures 7H-J, Supplemental Table 36**; *P <*0.0001).

A slightly different pattern emerges in the *Trh* neurons. As shown in **Figure 7K**, there is an apparent reduction in both *Nhlh2* and *Trh* signals in the DMH region of the hypothalamus of Snord116^del^ mice compared to WT mice. When comparing *Nhlh2* expression and localization in *Trh* neurons, total *Nhlh2* and nuclear-localized *Nhlh2* message is significantly reduced (*P =*0.0045 and *P =*0.0023, respectively), while cytoplasmic *Nhlh2* levels are similar between Snord116^del^ mice compared to WT mice (**Figures 7L-N, Supplemental Table 37**).

## Discussion

This study demonstrates a novel mechanism for *Snord116@* action in sequestering *Nhlh2* in the nuclear compartment, possibly protecting the message from degradation and allowing for increased Nhlh2 protein production. Using animal and cell-based models, the results show that *Snord116* co-expression increases nuclear *Nhlh2* expression levels in targeted regions of the brain as well as in a neuronal cell line. Furthermore, plasmid constructs containing the predicted *Snord116* binding site on the 3’UTR of *Nhlh2* are nuclear retained, while those lacking the binding site show no significant bias in the distribution of *Nhlh2* mRNA in cytoplasm and nuclei. These data confirm that *Nhlh2* is a target of *Snord116@* and suggest that some of the phenotypes seen in patients and mouse models of PWS may be due to altered expression patterns for *Nhlh2/NHLH2* mRNA and protein. Prior to these findings, the mechanism by which *Snord116@* functions had not been identified. The current study demonstrates a novel mechanism for *Snord116@* action in sequestering *Nhlh2* in the nuclear compartment, possibly protecting the message from degradation. This could allow for differential Nhlh2 protein translation under defined conditions, such as leptin stimulation or food deprivation. The sequestering of *Nhlh2* mRNA in the nuclear compartment of specific brain regions, especially the hypothalamus, helps clarify why we and others found no overall difference in *Nhlh2* expression level in WT versus Snord116^del^ or Snord116^m+/p-^ animals (i.e.^15^). It remains to be seen which other *Snord116@* targets are controlled in this way, and if the three major subgroups of *SNORD116* snoRNAs^8^ have different regulatory mechanisms.

Previous studies from our laboratory have demonstrated the presence of a sequence in the *Nhlh2* full-length mRNA 3’UTR (5’-UUUAGUCUUUUUCAGGGUUUUUUUGUG-3’) that is 100% conserved between human, chimp, macaque, rabbit, dog, mouse and rat *Nhlh2* mRNA sequences, and in 70 of 77 other animals examined^13^. Here we demonstrate that this sequence is necessary for *Nhlh2* mRNA partitioning in the nuclear compartment, as *Nhlh2-*myc with the partial 3’UTR does not respond to the presence of *Snord116* and shows no difference in nuclear versus cytoplasmic levels of *Nhlh2.* Indeed, our previous studies demonstrated that the shortened 3’UTR exhibits increased mRNA instability and produces less overall Nhlh2 protein^13^.

The finding that the presence of *Snord116@* results in nuclear localization of *Nhlh2* has several implications. First, while the data show that *Snord116* action requires the putative *Snord116-3* binding site in the 3’UTR, this could be an indirect function, or one not specifically requiring that binding site but another region within the full-length 3’UTR tail. The full-length mRNA for the human *NHLH2* gene is 2512 base pairs (2568 base pairs in mouse) with more than half of the sequence (1591 base pairs in human, and 1642 base pairs in mouse) making up the 3’UTR of the message. Interestingly, the putative *SNORD116* binding site in humans is found at the very end of the human *NHLH2* 3’UTR^13^. In the partial 3’UTR construct, the partial tail is missing the last 1154 nucleotides which contains the predicted *Snord116* binding motif, along with 20 AUUU motifs. The entire 3’UTR of *NHLH2* mRNA contains 80 AU-rich elements (AREs), according to a query done on ARESite2 database^24^. AREs have been implicated in modulating mRNA half-life and interacting with RNA-binding proteins. The RNA binding protein hnRNPU binds to and stabilizes the *Nhlh2* transcript, according to one report^25^. hnRNPU is expressed in the brain, where one study found that deficiency leads to chromatin restructuring, differential exon usage, and delayed neural commitment of progenitors from induced pluripotent stem cells^26^. HNRNPU is also called “Scaffold Attachment Factor” (SAF-A) and mutations in HNRNPU have been implicated in epilepsy, intellectual disability, developmental delay, and autism spectrum disorder^14,27–29^. Our laboratory showed that a human 3’UTR variant in NHLH2, rs11805084, contained within one of these AREs, changes the predicted 3D structure of the 3’UTR and destabilizes the mRNA^30^. While this particular variant is not near the putative *Snord116* binding site, these data support a role for mRNA stability in *Nhlh2* mRNA expression levels. Within the putative *SNORD116* binding site there are two predicted binding sites for the RNA binding protein ELAV1 (also known as HUR, data not shown, ARESite2 database^24^). ELAV1 is reported to change the subcellular distribution of mRNAs from the nucleus to the cytoplasm during immune stimulation and in this way protects its target mRNAs from degradation, until they are brought to the cytoplasm for translation^31^. While ELAV1 has never been shown to directly interact with snoRNAs, either directly, or within a regulatory pathway, this represents another area that warrants investigation.

The data in the current study support a role for the *Snord116* snoRNA in sequestering *Nhlh2* mRNA in the nucleus of specific neuron types, such as Pomc neurons, which we hypothesize helps to protect this mRNA from degradation over the long term. Emerging data from multiple studies suggest that this phenomenon called “bursty” mRNA transcription and translation^32–35^ may occur via long non-coding RNA Neat1, and RNA binding proteins^36–38^ in nuclear paraspeckles. In our studies, *Snord116@* is acting in a nuclear subdomain such as the nucleolus where it is known to reside^39^, to retain target mRNAs, protecting them from degradation prior to their release in a burst for protein translation. The mechanism and downstream signals that mediate the eventual release of target mRNAs for translation remain unknown and represent an important avenue for future studies examining this mRNA regulatory pathway.

In this study, partitioning of the *Nhlh2* mRNA between WT and Snord116^del^ mice was not as pronounced in Trh neurons as in Pomc neurons, although total *Nhlh2* mRNA levels were still reduced in the PWS mouse model. These results support previous findings, as well as our own, in which measurement of *Nhlh2* mRNA in whole hypothalamic RNA was not different between the two genotypes (i.e. ^40^). However, consistent with our data, *Nhlh2* mRNA is significantly reduced following single-cell sequencing of arcuate organoids from phenotypically normal or PWS induced pluripotent stem cells (personal communication, ^41^).

In summary, we have demonstrated co-localization of *Snord116@* and one of its mRNA target genes, *Nhlh2*, concurrently uncovering a possible mechanism for *Snord116* function in sequestering target mRNAs to protect them from degradation. As deletion of *SNORD116* snoRNAs represent the minimal genomic region associated with PWS, these findings have direct implications in the human condition of obesity and delayed pubertal development. Confirmation that *Nhlh2* is indeed a *Snord116@* target mRNA with its mRNA levels controlled by subcellular partitioning is a novel finding. The specific mechanism driving *Snord116-*mediated nuclear versus cytoplasmic localization may inform the development of new types of therapy to maintain high *Nhlh2* or other *Snord116@* target mRNAs in neurons, ultimately preserving regulatory pathways controlling body weight and reproductive phenotypes.

## Resource availability

### Materials availability

This study did not generate new unique reagents. RNAScope probes that were developed for this project are available from ACD Biotechne™. Requests for further information, resources, and reagents should be directed to, and will be fulfilled by, the lead contact, Deborah J. Good

### Data availability

Any additional information required to re-analyze the data reported in this paper is available from the lead contact, Deborah J. Good, upon request.

## Supporting information

Graphical Abstract

## Acknowledgments

The study was supported by grants from the Foundation for Prader-Willi Research to D.J.G. (#906446, 2022 and #532939, 2018), and the Margaret Cater Hepler Memorial Fund Summer Research Grant to S.A. We wish to thank the animal facility caretakers in the Integrated Sciences Building for their excellent care of the animals, and upkeep of the facility.

## Author contribution

Conceptualization by D.J.G and S.A.; Research design by D.J.G., S.A., and M.A.K.; D.J.G., S.A., and M.A.K. performed the *in vitro* and *in vivo* studies. C.K.T. consulted on confocal analysis. S.T. consulted on neuroanatomy. D.J.G. and S.A. wrote the first draft of the paper. All authors read and edited the final paper.

## STAR Methods

### Key Resources Table

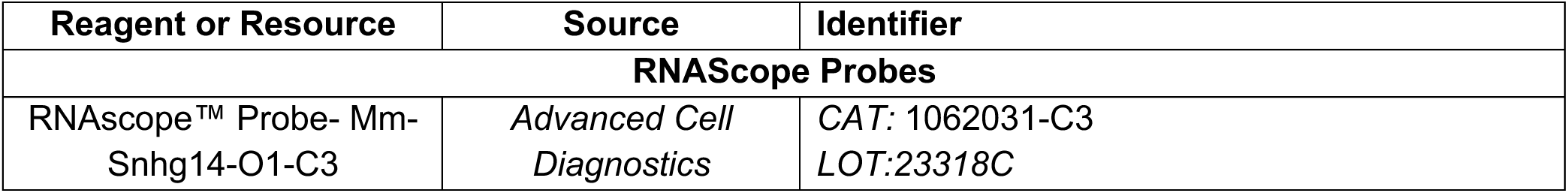
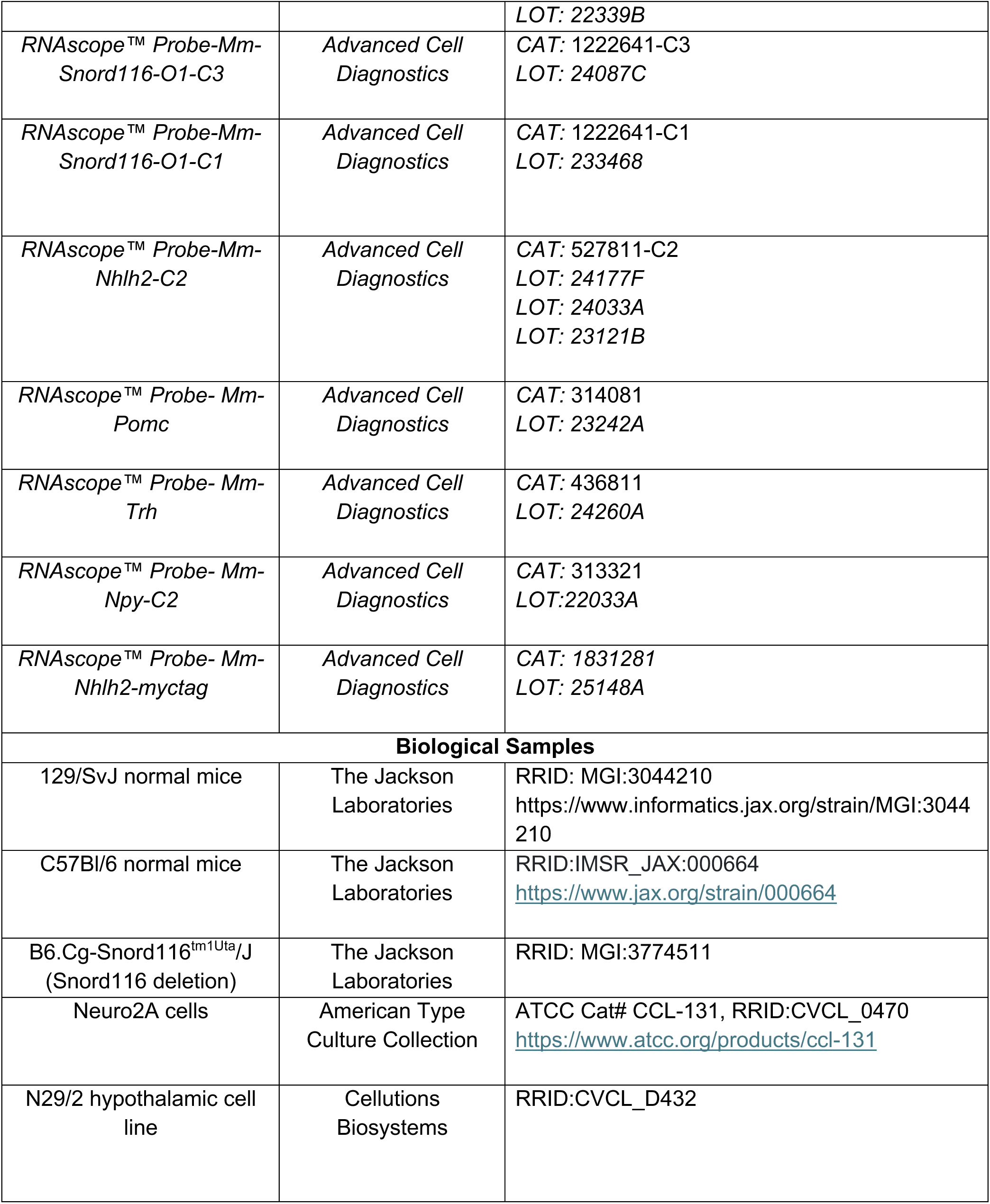
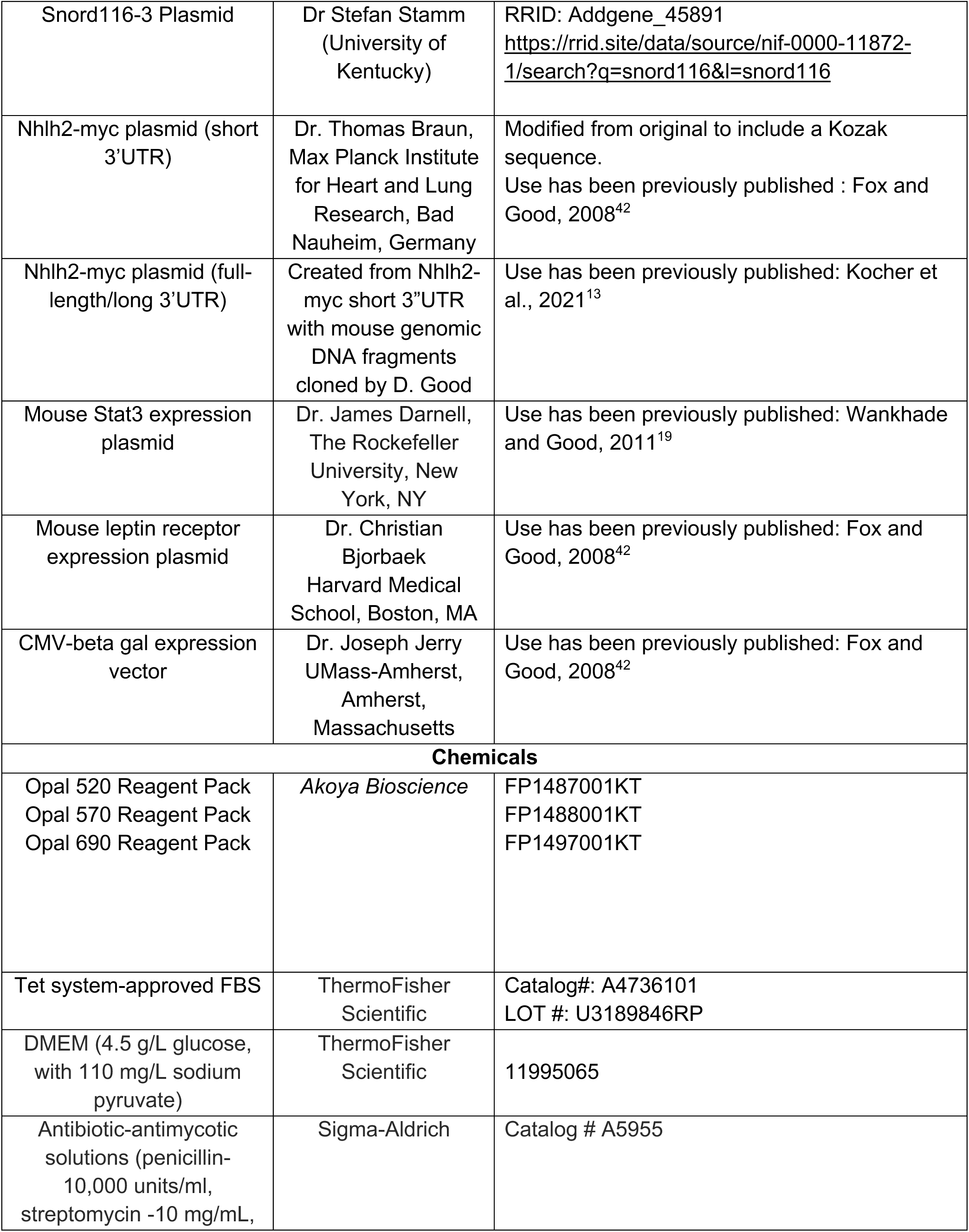
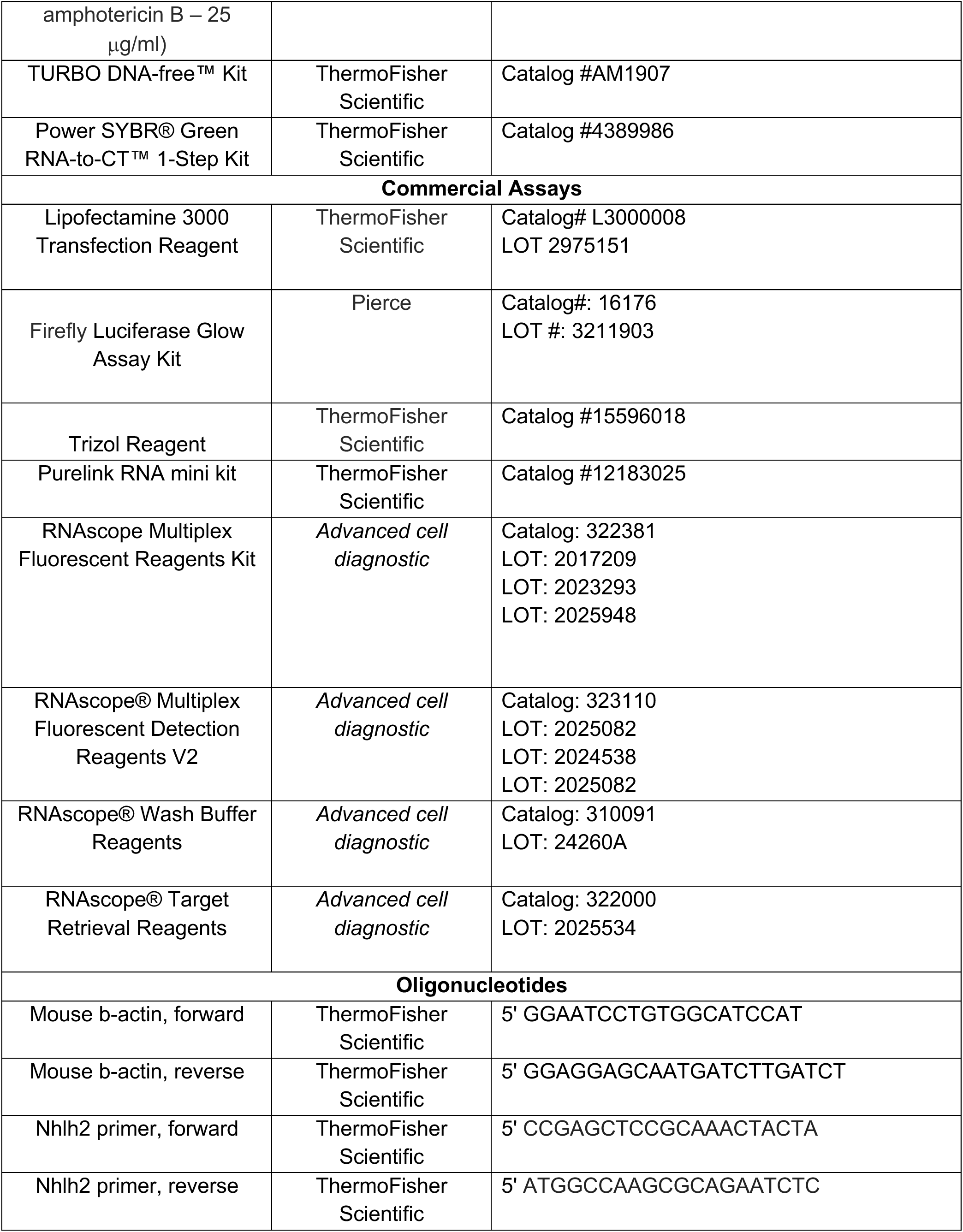
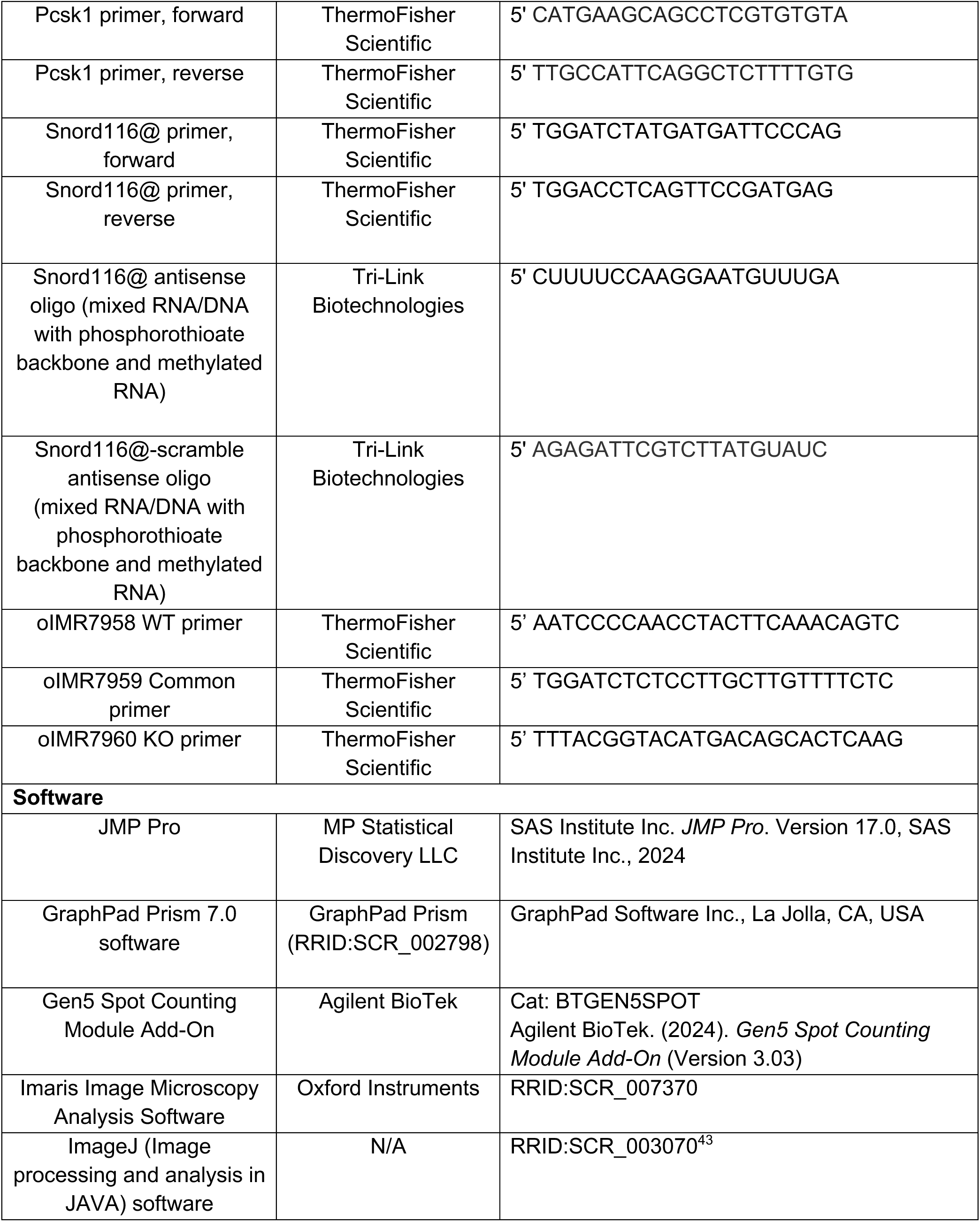

#### Experimental model and study participant details

##### Mice

Wildtype (WT) mice, strain 129Sv/J were used for the majority of the co-localization analyses. The B6.Cg-Snord116^tm1Uta^/J strain containing a deletion of the *Snord116* locus, which is on a C57Bl/6J strain (used as WT in comparative studies) were used in Figures 1 and 5. The 129Sv/J mice were bred using standard procedures with heterozygous male and female mice. Mice containing the *Snord116* deletion and age-matched C57Bl/6J WT mice were bred using two different schemas. Breeding pairs were set up between female heterozygous mice and WT males to generate heterozygous mice with the *Snord116* deletion coming from the maternal line. These animals carry the *Snord116* deletion on the maternal chromosome and do not have PWS. These were the mice used to set up the breeding mice that generated PWS offspring. Breeding mice to generate PWS mice were set up using heterozygous male mice, with WT females so that the *Snord116* deletion allele came from the paternal line. Mice from all strains were weaned at 3 weeks old and housed with littermates of the same sex. Genotyping was performed as reported^10,44^. All mice were held on a 7AM-7PM light cycle with standard Purina mouse chow (Purina diet 5001).

##### Cell Lines

The mouse neuroblastoma cell line, Neuro2A^45^, was maintained in T25 flasks in DMEM (1g/L glucose) and 10% fetal bovine serum (GE Healthcare #SH30396.03HI) with penicillin (50 units/mL) /streptomycin (50ug/mL) (Thermo # 15070063) at 4-6% CO2 and 37°C. Cells were detached from flasks using trypsin-EDTA (ThermoFisher # 25300054) and 6-well plates were seeded with 2 x 10^4^ - 10^5^ cells per well. Transfections were done 2-4 days post seeding using Opti-MEM® media (ThermoFisher # 31985070) and Lipofectamine® 3000 (ThermoFisher #L3000008) according to manufacturer’s instructions at 60-90% cell confluence.

The mouse hypothalamic cell line N29/2^14^ was cultured in T25 flasks using DMEM (4.5 g/L glucose, 110 mg/L sodium pyruvate; ThermoFisher, #11995065) supplemented with 10% fetal bovine serum (GE Healthcare, #SH30396.03HI) and penicillin (50 U/mL)/streptomycin (50 µg/mL) (ThermoFisher, #15070063). Cells were maintained at 37 °C in a humidified incubator with 4–6% CO₂ and used for experiments at passages 15 to 20. For passaging, Trypsin-EDTA (ThermoFisher, #25300054) was used to detach cells from the flask surface.

##### Tissue RNA isolation

For ad-lib fed conditions, mice were euthanized in home cages or separated from littermates if needed immediately before euthanizing. For 5-hour refeed conditions, between 5pm – 6pm mice were separated from home cages and singly housed in cages without food or bedding (to prevent mice from eating bedding). The following day, food was reintroduced at 9am until mice were euthanized at 2pm and tissue was collected.

For tissue dissection, mice were euthanized between 8 and 12 weeks old by CO2 asphyxiation and decapitated between 12pm and 2pm. Brains were then dissected using a Zivic-Miller mouse brain block and surgical tools, taking a hypothalamic slice between the optic chiasm, and the mammillary bodies. The slice was laid flat and dissected below the habenular region and on either side of the lateral ventricles to isolate a whole hypothalamus block. The block was lysed in TRIzol using a rotor stator homogenizer. TRIzol samples were stored at -20°C until RNA purification (1 week – 1 year). Fresh/autoclaved rotor stator homogenizers were used between samples to avoid cross-contamination.

Thawed TRIzol samples were purified using the TRIzol+Purelink RNA minikit (ThermoFisher #12183025) following manufacturer’s instructions for the TRIzol® Plus Total Transcriptome Isolation protocol. Purified RNA was then DNAse treated using TURBO DNA-free™ Kit (ThermoFisher #AM1907) according to manufacturer’s instructions, diluted to 60ng/uL in nuclease-free water, and stored at -80°C.

##### RT-QPCR

For RT-QPCR, Power SYBR® Green RNA-to-CT™ 1-Step Kit (ThermoFisher #4389986) was used according to manufacturer’s instructions. 10uL reactions were performed using 150nM final primer concentration. Primers were assessed for efficiency using a dilution series and fell within 90%-110% efficiency. 90ng RNA was used per 10uL reaction. Two to three technical replicates were performed. Control reactions for each sample (No reverse-transcriptase and no-template controls) were used for quality control. 384-well plates were run on the ViiA 7 Real-Time PCR System (ThermoFisher) according to RT-QPCR mix instructions and thermocycling conditions were not modified from suggested protocol (1-step annealing/extension at 60°C). Quality control measures including melt-curve analysis, technical replicate analysis, etc. were analyzed by thermocycler software and by operator; any major errors were excluded from analysis when appropriate, and/or new samples and plates were run when appropriate. Candidate reference genes for ddCT analysis were analyzed for appropriate reference controls. Mouse beta-actin was used as reference gene control for all experiments.

All RT-QPCR data were analyzed using Microsoft Excel 16 for Microsoft 365, and GraphPad Prism 9.0.0. Number of samples in statistical tests are described in respective figures. Error bars indicate ±SD. The 2^ddCT^ method of relative quantification was used. Statistical significance tests performed on respective ddCT values from which Relative Quantification values are derived. Significance is expressed at *p<0.05, **p<0.01, ***p<0.001. A two-way ANOVA with Bonferroni correction was used for relative expression of RNA normalized to WT or control conditions.

##### Tissue Preparation for In situ Hybridization (RNAScope™)

The trans cardiac perfusion method was used to remove blood cells from tissues and maintain brain integrity for the *in-situ* hybridization. Between 12-3 PM, ad lib-fed mice were rendered unconscious using CO_2_ asphyxiation, and consciousness confirmed by a non-responsive toe pinch. The chest cavity was then opened, and penetration to the left ventricle was done while the heart was still beating using a 25-gauge infusion blunted needle. After cutting the atrium, phosphate buffered saline (PBS) was pumped into the ventricle using a perfusion pulsating pump (BS-900 -0G, Braintree Scientific, Braintree, MA).). The pressure was set to .72 ml/min with 1/16” ID tubing. Perfusion of ∼20 ml of PBS was continued until the liver was noticeably pale, approximately 25 minutes. Perfusion with 4% paraformaldehyde (PFA) in PBS was then started by switching the solutions and continued with the same pressure until the tail became noticeably stiff (approximately 20 minutes). The brain was then removed from the cranial cavity, and dissected into anterior, medial, posterior sections coronally using the Zivic-Miller mouse brain matrix. Each section was post-fixed in 4% PFA for 24 hours at 4^0^C and then held at 4^0^C in 70% ethanol until embedding. All tissues were sent to the Virginia-Maryland Veterinary Medicine Histology Core Facility at Virginia Tech for further processing and embedding into paraffin wax.

##### Sectioning and Staining Histology

Embedded blocks were cut with the 10μm thickness using a tissue microtome and placed in a 40^0^C water bath prior to placement on glass slides (VWR® Superfrost® Plus Micro Slide, VWR International, Radnor, PA, USA Cat. No. 48311-703). Slides were placed on a slide warmer for 24 hours to dry and then stored at room temperature prior to histological staining or the RNAscope assay. To determine location of specific brain regions within the coronal sections, every 10^th^ slide was subject to Hematoxylin and Eosin (H&E) or Methyl Green staining.

##### In situ Hybridization

*In situ* hybridization studies were performed using the RNAscope® method. (Bio-techne/Advanced Cell Diagnostics, Newark, CA). This method is an advanced platform for *in situ* RNA detection and was used in this study to help preserve tissue morphology and architecture and allowing multiplex analysis of RNA expression with high sensitivity and specificity. The manufacturers’ procedure was used for all steps, with the target retrieval and protease steps optimized to the tissues and probes being used. A positive and negative probe set was used to initially perform optimization with individual tissue types. Opal dye sets (Akoya Biosciences, Marbough, MA) were used as the fluorophors, and slides were counterstained with DAPIO blue stain (Bio-techne/Advanced Cell Diagnostics, Newark, CA).

##### Imaging

Two different methods of imaging were used for all the studies. First, an inverted fluorescent microscope (Cytation 5 imaging multimode reader from Agilent biotechnology, Santa Clara, CA) capable of magnifications at 2.5, 4x, 20x, and 60x was used to initially visualize the tissues. After initial imaging and analysis, slides were viewed using Leica SP8 (Leica Microsystems GmbH, Wetzlar, Germany), which allowed for more accurate and 3D visualization using 63x magnification microscope with different zoom options. Confocal image datasets were processed and analyzed using Imaris™ software (Oxford Instruments, version 10.1; RRID: SCR_007370). Z-stack images were first imported into Imaris in their native Leica lif format, with voxel size and step interval retained from microscope acquisition metadata and based on fluorescence intensity thresholds. For 3D visualization, the Surpass mode in Imaris™ was used. Z-stack image volumes were rendered into three-dimensional reconstructions using the “Volume” modules. To assess colocalization, the Imaris™ Coloc module was used along with manual thresholding.

##### Signal Quantification

The Cytation 5 microscope is also equipped with the Gen% spot counting module that facilitates quantification of different puncta, allowing for simultaneous quantification of different signals within the same cell. The software provides the chance to simultaneously count individual dots inside and outside of the neuron’s nucleus and quantify the number of Nhlh2 and Snhg14/Snord116 puncta. This was done in a non-blinded way by the operator. To count cytoplasmic signals, the total amount of detected signals inside and outside of the neurons were deducted from the nucleus. Signals found on the border of the DAPI stained nucleus were considered as cytoplasmic. All analyses were performed on raw, unprocessed images without any brightness or contrast adjustments to ensure objective quantification. Nuclei were identified based on DAPI staining using predefined size (5–100 µm diameter) and intensity thresholds. Within each defined nuclear region, fluorescent spots corresponding to RNA signals were detected using spot size parameters of 0.5–5 µm for Nhlh2 and 1–5 µm for Snord116/Snhg14, with optimized intensity thresholds to distinguish true signal from background fluorescence. Spot counts per cell were automatically calculated and exported for statistical analysis. Each dataset was subsequently reviewed by the researcher to identify and correct any errors or mismatches introduced by the automated detection. The software’s advanced segmentation features, including object separation and edge-detection tools, were enabled to enhance spot detection accuracy.

##### Cell transfection

For *Snord116* knockdown, anti-sense oligos (ASO) were 20nt in the structure of 5nt-10nt-5nt RNA/DNA/RNA with phosphorothioate backbone and methylated RNA.Snord116-targetting ASOs and control ASOs targeting nothing were transfected into Neuro2A cells to a final concentration of 100nM, as optimized for *Snord116* knockdown. For cell culture, 24 hours post transfection, cell culture media was removed, and cells were lysed with 1mL of TRIzol® Reagent (ThermoFisher #15596018) directly in 6-well culture plates. TRIzol samples were frozen at -20°C in microfuge tubes until purified. RNA and RT-QPCR procedures were described above.

For transfection experiments using N29/2 cells, cells were seeded into either 6-well plates or 2-well Nunc® Lab-Tek® Chamber Slide™ systems (ThermoFisher Scientific, Waltham, MA, USA; #41122600) for in situ hybridization applications, depending on the experimental requirements described below. Cells were seeded at a density of 6 × 10^5^ cells per chamber, based on the Lipofectamine® 3000 transfection protocol (ThermoFisher, #L3000008). Transfection was performed 2–4 days post-seeding using Opti-MEM® reduced serum medium (ThermoFisher, #31985070), following the manufacturer’s instructions, when cultures reached 60–90% confluence, plus transfection-certified tet-minus fetal bovine serum. A total of 800 ng of plasmid DNA per chamber was used, consisting of 200 ng each of Leptin receptor, Stat3, ±Snord116, and Nhlh2 full-length (Myc-tagged) expression vectors. Leptin receptor and Stat3 constructs were included to ensure co-expression of key regulatory components of Nhlh2, in alignment with prior studies. Nhlh2 – full length, short-tail and human SNV variant constructs were used in parallel. RNAscope® in situ hybridization was performed between 24- and 72- hours post-transfection, following the manufacturer’s protocol for adherent cell assays. Transfections were performed in triplicate, with total input DNA kept constant across conditions by supplementing with empty vector where necessary.

##### Statistical Analysis

In JMP, *Nhlh2* expression was treated as the dependent variable and modeled as a function of *Snhg14* expression using the Generalized Linear Model (GLM) platform. The distribution was set to Negative Binomial, and a log link function was applied. The overdispersion parameter was estimated from the data. For each region, we extracted regression coefficients, standard errors, 95% confidence intervals, Wald chi-square statistics, and p-values. These model outputs were used to assess the direction and significance of the relationship between *Snhg14, Snord116*, and *Nhlh2* expression. All figures were generated using custom Python scripts (Python 3.11, Matplotlib 3.7) to ensure consistent formatting and visual clarity across brain regions.

## Supplemental Information

