## Supplemental Tables for "Subcellular partitioning of *Nhlh2* mRNA reveals how *SNORD116* loss contributes to Prader-Willi Syndrome"

**Supplemental Table 1: Summary of Mann–Whitney U Tests for *Nhlh2* full-length 3’ UTR levels among two groups of *Snord116* + and *Snord116*– (N29/2 hypothalamic neurons)—Figure 2E**

| ****Expression Compartment**** | Snord116 + ****Median (n = 34)**** | Snord116– ****Median (n = 37)**** | ****Test**** | ****U Statistic**** | ****P Value**** | ****Summary**** | ****Significant?**** |
| --- | --- | --- | --- | --- | --- | --- | --- |
| ****Total**** | 0.000 | 0.000 | Mann–Whitney U | 484 | 0.0497 | * | Yes |
| ****Nuclear**** | 0.000 | 0.000 | Mann–Whitney U | 420.5 | 0.0007 | *** | Yes |
| ****Cytoplasmic**** | 0.000 | 0.000 | Mann–Whitney U | 628 | 0.9888 | ns | No |
| ns: Not significant  *: Significant at p < 0.05  **: Significant at p < 0.01  ***: Significant at p < 0.001 | | | | | | | |

**Supplemental Table 2: Summary of Mann–Whitney U Tests for *Nhlh2* partial 3’ UTR levels among two groups of *Snord116* + and *Snord116* – (N29/2 hypothalamic neurons)**—**Figure 2F**

| Expression Compartment | Snord116⁺ Median (n = 45) | Snord116⁻ Median (n = 24) | Statistical Test | P Value | Summary | Significant (P < 0.05)? |
| --- | --- | --- | --- | --- | --- | --- |
| Total | 1.000 | 1.000 | Mann–Whitney U = 463 | 0.2926 | ns | No |
| Nuclear | 0.000 | 0.000 | Mann–Whitney U = 418 | 0.0546 | ns | No |
| Cytoplasmic | 1.000 | 1.000 | Mann–Whitney U = 536 | 0.9640 | ns | No |
| ns: Not significant  *: Significant at p < 0.05  **: Significant at p < 0.01  ***: Significant at p < 0.001 | | | | | | |

**Supplemental Table 3: Coefficient Estimates from Negative Binomial Regression in Dentate Gyrus—Figure 4A, Supplemental Figure 1A**

|  | Estimate | Std Error | Wald X2 | P-value | 95% CI |
| --- | --- | --- | --- | --- | --- |
| Intercept | –0.1661 | 0.0993 | 2.80 | 0.0944 | [-0.3606, 0.0285] |
| *Snhg14 expression* | 0.5771 | 0.0696 | 68.71 | <0.0001 | [0.4406, 07135] |
| ns: Not significant  *: Significant at p < 0.05  **: Significant at p < 0.01  ***: Significant at p < 0.001 | | | | | |

**Supplemental Table 4. Unit Incidence Rate Ratio for** ***Snhg14* Predicting *Nhlh2* Expression in Dentate Gyrus—Figure 4A Supplemental Figure 1A**

|  | IRR | 95% CI |
| --- | --- | --- |
| Snhg14 expression | 1.781 | [1.554, 2.041] |
| Snhg14 expression | 10.058 | [5.827, 17.360] |

**Supplemental Table 5. Coefficient Estimates from Negative Binomial Regression in medial habenula—Figure 4D,**

**Supplemental Figure 1B**

|  | **Estimate** | **Std Error** | **Wald X²** | **p-value** | **95% CI** |
| --- | --- | --- | --- | --- | --- |
| **Intercept** | 0.099 | 0.071 | 1.94 | 0.164 | [–0.040, 0.237] |
| ***Snhg14* expression** | 0.601 | 0.050 | 143.95 | <0.0001 | [0.503, 0.699] |
| ns: Not significant  *: Significant at p < 0.05  **: Significant at p < 0.01  ***: Significant at p < 0.001 | | | | | |

**Supplemental Table 6. Incidence Rate Ratios for**  ***Snhg14***  **Expression Predicting *Nhlh2* in the medial habenula—Figure 4D,** **Supplemental Figure 1B**

|  | IRR | 95% CI |
| --- | --- | --- |
| Snhg14 expression | 1.824 | [1.653, 2.012] |
|  | **IRR** | **95% CI** |
| Snhg14 expression | 11.07 | [7.47, 16.39] |

**Supplemental Table 7:** **Coefficient Estimates from Negative Binomial Regression cortex**—**Figure 4G, Supplemental Figure 1C**

|  | Estimate (β) | Std. Error | Wald χ² | df | p-value | 95% CI (β) |
| --- | --- | --- | --- | --- | --- | --- |
| Intercept | –1.238 | 0.205 | 36.50 | 1 | < .0001 | [–1.640, –0.837] |
| *Snord116*/nucleus | +1.088 | 0.221 | 24.27 | 1 | < .0001 | [+0.655, +1.520] |
| ns: Not significant  *: Significant at p < 0.05  **: Significant at p < 0.01  ***: Significant at p < 0.001 | | | | | | |

**Supplemental Table 8. Incidence Rate Ratios (IRRs) for *Snord116* Expression Predicting *Nhlh2* per Nucleus in Cortex—Figure 4,** **Supplemental Figure 1C**

|  | **IRR** | **95% CI (IRR)** | **Notes** |
| --- | --- | --- | --- |
| ***Snord116*/nucleus** | 2.967 | [1.925, 4.573] | Per unit increase in Snord116/nucleus |
| ***Snord116*/nucleus** | 8.802 | [3.705, 20.913] | Across full range |

**Supplemental Table 9. Coefficient Estimates from Negative Binomial Regression in the Thalamus**—**Figure 4J,**

**Supplemental Figure 1. D**

|  | **Estimate** | **Std Error** | **Wald X²** | **p-value** | **95% CI** |
| --- | --- | --- | --- | --- | --- |
| **Intercept** | –1.238 | 0.205 | 36.50 | <0.0001 | [–1.640, –0.837] |
| ***Snord116*** | 1.088 | 0.221 | 24.27 | <0.0001 | [0.655, 1.520] |
| ns: Not significant  *: Significant at p < 0.05  **: Significant at p < 0.01  ***: Significant at p < 0.001 | | | | | |

**Supplemental Table 10. Incidence Rate Ratios for *Snord116* Predicting *Nhlh2* Expression in the Thalamus—Figure 4J, Supplemental Figure 1D**

|  | **IRR** | **95% CI** |
| --- | --- | --- |
| **Unit incident rate ratio** |  |  |
| ***Snord116*** | 2.97 | [1.92, 4.57] |
| **Full range of expression** |  |  |
|  | IRR | 95% CI |
| ***Snord116*** | 8.80 | [3.70, 20.91] |

**Supplemental Table 11: Coefficient Estimates from Negative Binomial Regression in the Paraventricular Nucleus of Hypothalamus—Figure 4 M-O, Supplemental Figure 1F**

|  | **Estimate** | **Std. Error** | **Wald X^2^** | **P-value** | **95% Confidence Interval** |
| --- | --- | --- | --- | --- | --- |
| **Intercept** | 1.498 | 0.112 | 178.33 | <0.0001 | [1.278, 1.718] |
| ***Snhg14* expression** | 0.115 | 0.028 | 16.59 | <0.0001 | [0.060, 0.170] |
| ns: Not significant  *: Significant at p < 0.05  **: Significant at p < 0.01  ***: Significant at p < 0.001 | | | | | |

**Supplemental Table 12. Incidence Rate Ratios for *Snhg14* Predicting *Nhlh2* Expression in the Paraventricular Nucleus of Hypothalamus—Figure 4M-O, Supplemental Figure 1F**

|  | IRR | 95% CI |
| --- | --- | --- |
| Unit Incidence Ratio |  |  |
| *Snhg14* | 1.12 | [1.061,1.185] |
| Full range of expression |  |  |
|  | **IRR** | **95% CI** |
| *Snhg14* | 3.16 | [1.816, 5.499] |

**Supplemental Table 13: Parameter Estimates from Negative Binomial Regression Predicting *Nhlh2* Expression per Nucleus in arcuate nucleus of hypothalamus**—**Figure 4N,** **Supplemental Figure 1E**

|  | Estimate | Std Error | Wald X2 | P-value | 95% CI |
| --- | --- | --- | --- | --- | --- |
| Intercept | -1.095 | 0.164 | 44.76 | <.0001 | [-1.415, -0.774] |
| Snhg14 expression | 0.464 | 0.123 | 14.25 | 0.0002 | [0.223, 0.705] |

**Supplemental Table 14**: **Incidence Rate Ratio (IRR) for *Snhg14* Predicting *Nhlh2* Expression in Arcuate Nucleus of Hypothalamus**—**Figure 4N, Supplemental Figure 1E**

|  | IRR | 95% CI |
| --- | --- | --- |
| Unit Incidence Ratio |  |  |
| *Snhg14* | 1.591 | [1.250,2.024] |
| Full range of expression |  |  |
|  | **IRR** | **95% CI** |
| *Snhg14* | 16.21 | [3.82, 68.83] |

**Supplemental Table 15 : Mann–Whitney U analysis of overlap volume ratio between two** **fluorescent channels—Figure 5L**

| Metric | Value |
| --- | --- |
| Statistical test | Mann–Whitney U test (two-tailed, exact) |
| U statistic | 73 |
| Sum of ranks (Coloc + / Coloc −) | 881 / 2774 |
| p-value | < 0.0001 |
| Significance | **** (p < 0.0001) |
| Median overlap (Coloc +) | 0.030 (n = 12 neurons) |
| Median overlap (Coloc −) | 0.000 (n = 73 neurons) |
| Median difference (Hodges–Lehmann) | −0.030 |
| Median difference (actual) | −0.030 |

**Supplemental Table 16 : Mann–Whitney U analysis of nuclear volume between co-expression –positive and –negative neurons—Figure 5M**

| Metric | Value |
| --- | --- |
| Statistical test | Mann–Whitney U test (two-tailed, exact) |
| U statistic | 190.5 |
| Sum of ranks (Coloc + / Coloc −) | 763.5 / 2892 |
| p-value | 0.0012 |
| Significance | ** (p < 0.01) |
| Median nuclear volume (Coloc +) | 68.15 (n = 12 neurons) |
| Median nuclear volume (Coloc −) | 6.13 (n = 73 neurons) |
| Median difference (Hodges–Lehmann) | −54.90 |
| Median difference (actual) | −62.02 |

**Supplemental Table 17: Chi-Square Analysis of *Nhlh2* Subcellular Localization by *Snhg14* Expression in Pomc Neurons—Figure 5N**

| ****Parameter**** | ****Result**** |
| --- | --- |
| Statistical test | Fisher’s exact test |
| P value | < 0.0001 |
| Significance | **** |
| Interpretation | Significant |
| Table dimensions | 4 rows × 2 columns |

**Supplemental Table 18: Distribution of *Nhlh2* Subcellular Localization *in Snhg14+* and *Snhg14*− Pomc Neurons—Figure 5N**

| Nhlh2 Localization | Snhg14+ | Snhg14− | Total | % Expressed (column) | % Not Expressed (column) |
| --- | --- | --- | --- | --- | --- |
| None | 2 | 4 | 6 | 3.39% | 8.33% |
| Cytoplasmic | 3 | 21 | 24 | 5.08% | 43.75% |
| Nuclear | 3 | 3 | 6 | 5.08% | 6.25% |
| Both partitions | 51 | 20 | 71 | 86.44% | 41.67% |

**Supplemental Table 19: Summary of Fisher’s Exact Test for “None” *Nhlh2* Localization across *Snhg14* Groups—Figure 5N**

| Parameter | Value |
| --- | --- |
| Test | Fisher’s exact test |
| P value | 0.4048 |
| Significance | ns |
| Two-sided? | Yes |
| Significant (P < 0.05)? | No |

**Supplemental Table 20: Fisher’s Exact Test for Cytoplasmic *Nhlh2* Localization across *Snhg14* Groups—Figure 5N**

| ****Parameter**** | ****Value**** |
| --- | --- |
| ****Test**** | Fisher’s exact test |
| ****P value**** | < 0.0001 |
| ****Significance**** | **** |
| ****Two-sided?**** | Yes |
| ****Significant (P < 0.05)?**** | Yes |
| ****Table Dimensions**** | 2 × 2 |

**Supplemental Table 21. Fisher’s Exact Test for Nuclear *Nhlh2* Localization across *Snhg14* Groups—Figure 5N**

| Parameter | Value |
| --- | --- |
| Test | Fisher’s exact test |
| P value | > 0.9999 |
| Significance | ns |
| Two-sided? | Yes |
| Significant (P < 0.05)? | No |
| Table Dimensions | 2 × 2 |

### **Supplemental Table 22. Fisher’s Exact Test for Dual (Both) *Nhlh2* Localization across *Snhg14* Groups—Figure 5N**

| Parameter | Value |
| --- | --- |
| Test | Fisher’s exact test |
| P value | < 0.0001 |
| Significance | **** |
| Two-sided? | Yes |
| Significant (P < 0.05)? | Yes |
| Table Dimensions | 2 × 2 |

**Supplemental Table 23. Mann–Whitney U analysis of Total, Nuclear, and Cytoplasmic *Nhlh2* Expression in *Snhg14⁺* vs. *Snhg14⁻* Pomc Neuron—Figure 5O**

| Expression Compartment | Test Used | Median (Snhg14⁺) | Median (Snhg14⁻) | U Statistic | P Value | Significance |
| --- | --- | --- | --- | --- | --- | --- |
| ****Total**** | Mann–Whitney Test | 6.0 (n = 59) | 3.0 (n = 48) | 812.5 | 0.0001 | *** |
| ****Nuclear**** | Mann–Whitney Test | 3.0 (n = 59) | 0.0 (n = 48) | 561 | < 0.0001 | **** |
| ****Cytoplasmic**** | Mann–Whitney Test | 3.0 (n = 59) | 2.0 (n = 48) | 1247 | 0.2830 | ns |
| ns: Not significant  *: Significant at p < 0.05  **: Significant at p < 0.01  ***: Significant at p < 0.001 | | | | | | |

**Supplemental Table 24: Chi-Square Analysis of *Nhlh2* Subcellular Localization by Snhg14 Expression in Trh Neurons—Figure 6E**

| ****Parameter**** | ****Value**** |
| --- | --- |
| ****Test**** | Fisher’s exact test |
| ****P value**** | 0.0361 |
| ****Significance**** | * |
| ****Two-sided?**** | Yes |
| ****Significant (P < 0.05)?**** | Yes |
| ****Table Dimensions**** | 4 × 2 |

**Supplemental Table 25: Contingency Analysis of *Snhg14* Expression by *Nhlh2* Localization in Trh Neurons—Figure 6E.**

| *Nhlh2* Localization | Snhg14+ | Snhg14− | Total | % Expressed (column) | % Not Expressed (column) |
| --- | --- | --- | --- | --- | --- |
| None | 1 | 3 | 4 | 2.13% | 11.54% |
| Cytoplasmic | 3 | 6 | 9 | 6.38% | 23.08% |
| Nuclear | 6 | 1 | 7 | 12.77% | 3.85% |
| Both partitions | 37 | 16 | 53 | 78.72% | 61.54% |

**Supplemental Table 26: Summary of Fisher’s Exact Test for “None” *Nhlh2* Localization Across *Snhg14* Groups—Figure 6E**

| ****Parameter**** | ****Value**** |
| --- | --- |
| ****Test**** | Fisher’s exact test |
| ****P value**** | 0.1260 |
| ****Significance**** | ns |
| ****Two-sided?**** | Yes |
| ****Significant (P < 0.05)?**** | No |
| ****Table Dimensions**** | 2 × 2 |

**Supplemental Table 27: Summary of Fisher’s Exact Test for “Cytoplasmic” *Nhlh2* Localization Across *Snhg14* Groups—Figure 6E**

| ****Parameter**** | ****Value**** |
| --- | --- |
| ****Test**** | Fisher’s exact test |
| ****P value**** | 0.0606 |
| ****Significance**** | ns |
| ****Two-sided?**** | Yes |
| ****Significant (P < 0.05)?**** | No |
| ****Table Dimensions**** | 2 × 2 |

**Supplemental Table 28: Summary of Fisher’s Exact Test for “Nuclear” *Nhlh2* Localization Across *Snhg14* Groups—Figure 6E**

| ****Parameter**** | ****Value**** |
| --- | --- |
| ****Test**** | Fisher’s exact test |
| ****P value**** | 0.4094 |
| ****Significance**** | ns |
| ****Two-sided?**** | Yes |
| ****Significant (P < 0.05)?**** | No |
| ****Table Dimensions**** | 2 × 2 |

**Supplemental Table 29: Summary of Fisher’s Exact Test for “Dual (Both) *Nhlh2* Localization Across *Snhg14* Groups—Figure 6E**

| ****Parameter**** | ****Value**** |
| --- | --- |
| ****Test**** | Fisher’s exact test |
| ****P value**** | 0.1701 |
| ****Significance**** | ns |
| ****Two-sided?**** | Yes |
| ****Significant (P < 0.05)?**** | No |
| ****Table Dimensions**** | 2 × 2 |

**Supplemental Table 30: Mann–Whitney U analysis of Trh Expression Between *Snhg14⁺* and *Snhg14⁻* Cells Across Subcellular Compartments—Figure 6F**

| ****Expression Compartment**** | ****Test**** | ****Snhg14⁻ Median (n)**** | ****Snhg14⁺ Median (n)**** | ****Difference (Actual / HL)**** | ****P Value**** | ****P Value Summary**** |
| --- | --- | --- | --- | --- | --- | --- |
| ****Total**** | Mann–Whitney U | 3.5 (n = 26) | 6.5 (n = 48) | −3.0 / −2.0 | 0.0299 | * |
| ****Nuclear**** | Mann–Whitney U | 2.0 (n = 26) | 3.0 (n = 48) | −1.0 / 0.0 | 0.6085 | ns |
| ****Cytoplasmic**** | Mann–Whitney U | 1.0 (n = 26) | 4.0 (n = 48) | −3.0 / −2.0 | 0.0003 | *** |
| ns: Not significant  *: Significant at p < 0.05  **: Significant at p < 0.01  ***: Significant at p < 0.001 | | | | | | |

**Supplemental Table 31: Distribution of *Snhg14*  across genotypes in LH—Figure 7B**

| Genotype | Snhg14⁺ | Snhg14⁻ | Total | ****Snhg14⁺ (%)**** | ****Snhg14⁻ (%)**** |
| --- | --- | --- | --- | --- | --- |
| WT | 37 | 63 | 100 | 52.86% | 48.46% |
| Snord116del | 33 | 67 | 100 | 47.14% | 51.54% |
| **Total** | **70** | **130** | **200** | **100%** | **100%** |

**Supplemental Table 32: Fisher’s Exact Test and Effect Size Summary for *Snhg14* Expression Across Genotypes in LH—Figure 7B**

| ****Category**** | ****Result**** |
| --- | --- |
| ****Test Used**** | Fisher’s exact test |
| ****P value**** | 0.6567 |
| ****P value summary**** | ns |
| ****One- or two-sided?**** | Two-sided |
| ****Significant (P < 0.05)?**** | No |

**Supplemental Table 33: Distribution of *Snhg14* *–Nhlh2* co-expression by genotype in LH—Figure 7C**

| Genotype | Nhlh2 Co-expression (%) | No Nhlh2 co-expression (%) | n (neurons) |
| --- | --- | --- | --- |
| WT | 40.7% | 59.3% | 189 |
| Snord116⁻/⁻ | 22.6% | 77.4% | 186 |

**Supplemental Table 34: Chi-square analysis of co-expression of *Snhg14* with *Nhlh2* in WT and Snord116^del^ in LH—Figure 7C**

| Metric | Value |
| --- | --- |
| Statistical test | Chi-square test with Yates’ correction |
| χ² (df) | 13.45 (1) |
| z-score | 3.67 |
| p-value | 0.0002 |
| Significance | *** (p < 0.001) |
| Attributable risk (WT − Snord116⁻/⁻) | 0.182 |
| 95% CI (attributable risk) | 0.084 – 0.275 |
| Number needed to treat (NNT) | 5.51 |
| 95% CI (NNT) | 3.64 – 11.92 |

**Supplemental Table 35. Summary of Mann–Whitney U Tests for *Nhlh2* Counts in Lateral hypothalamus neruons (WT vs. Snord116^del^)—Figure 7D-F**

| ****Expression Compartment**** | ****Test**** | ****WT Median (n)**** | ****Snord116del Median (n)**** | ****Difference (Actual / HL)**** | ****U Statistic**** | ****P Value**** | ****Significance**** |
| --- | --- | --- | --- | --- | --- | --- | --- |
| ****Total**** | Mann–Whitney U | 2.0 (n = 189) | 1.0 (n = 186) | −1.0 / −1.0 | 8038 | < 0.0001 | **** |
| ****Cytoplasmic**** | Mann–Whitney U | 1.0 (n = 79) | 0.0 (n = 186) | −1.0 / −1.0 | 3094 | < 0.0001 | **** |
| ****Nuclear**** | Mann–Whitney U | 1.0 (n = 189) | 0.0 (n = 186) | −1.0 / −1.0 | 9588 | < 0.0001 | **** |
| ns: Not significant  *: p < 0.05  **: p < 0.01  ***: p < 0.001  ****: p < 0.0001 | | | | | | | |

**Supplemental Table 36. Summary of Mann–Whitney U Tests for *Nhlh2* Counts in Pomc neruons (WT vs. Snord116^del^) Figure 7: H-J**

| ****Expression Compartment**** | ****Test**** | ****WT Median (n)**** | ****Snord116del Median (n)**** | ****Difference (Actual / HL)**** | ****U Statistic**** | ****P Value**** | ****Significance**** |
| --- | --- | --- | --- | --- | --- | --- | --- |
| ****Total**** | Mann–Whitney U | 7.0 (n = 84) | 3.0 (n = 77) | −4.0 / −4.0 | 1182 | < 0.0001 | **** |
| ****Cytoplasmic**** | Mann–Whitney U | 3.0 (n = 84) | 1.0 (n = 77) | −2.0 / −2.0 | 1812 | < 0.0001 | **** |
| ****Nuclear**** | Mann–Whitney U | 3.0 (n = 84) | 1.0 (n = 77) | −2.0 / −2.0 | 1403 | < 0.0001 | **** |
| ns: Not significant  *: p < 0.05  **: p < 0.01  ***: p < 0.001  ****: p < 0.0001 | | | | | | | |

**Supplemental Table 37. Summary of Mann–Whitney U Tests for *Nhlh2* Counts in Trh neruons (WT vs. Snord116^del^) Figure 7 L-N**

| ****Expression Compartment**** | ****Test**** | ****WT Median (n)**** | ****Snord116del Median (n)**** | ****Difference (Actual / HL)**** | ****U Statistic**** | ****P Value**** | ****Significance**** |
| --- | --- | --- | --- | --- | --- | --- | --- |
| ****Total**** | Mann–Whitney U | 6.0 (n = 36) | 3.0 (n = 41) | −3.0 / −2.0 | 463.5 | 0.0045 | ** |
| ****Nuclear**** | Mann–Whitney U | 3.0 (n = 36) | 2.0 (n = 41) | −1.0 / −1.0 | 447.5 | 0.0023 | ** |
| ****Cytoplasmic**** | Mann–Whitney U | 2.0 (n = 36) | 1.0 (n = 41) | −1.0 / 0.0 | 673.5 | 0.5051 | ns |
| ns: Not significant  *: p < 0.05  **: p < 0.01  ***: p < 0.001  ****: p < 0.0001 | | | | | | | |
