## Supplementary figures and images for "Subcellular partitioning of *Nhlh2* mRNA reveals how *SNORD116* loss contributes to Prader-Willi Syndrome"

### Graphical Abstract

Graphical Abstract
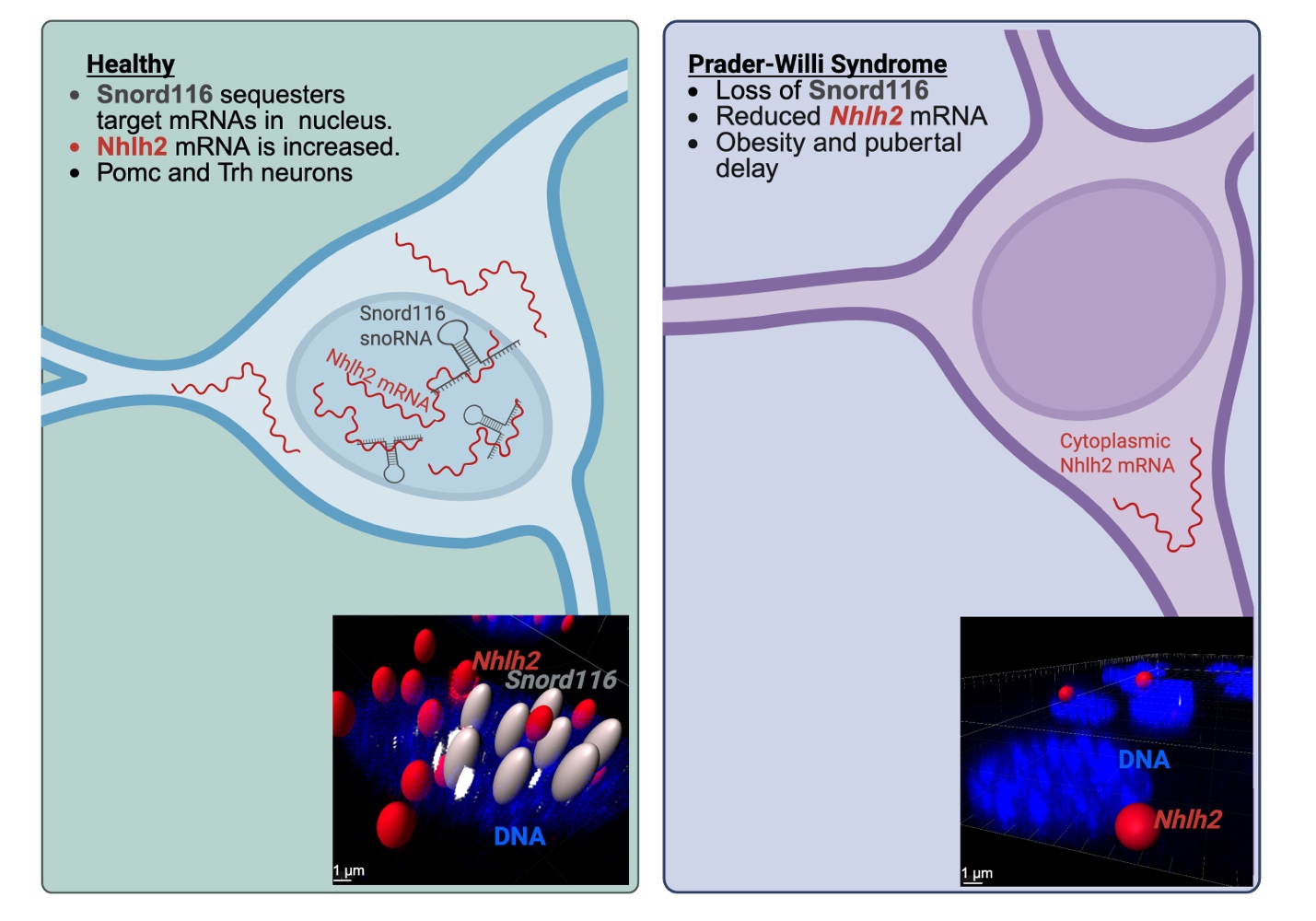

### Supplemental Figure 1

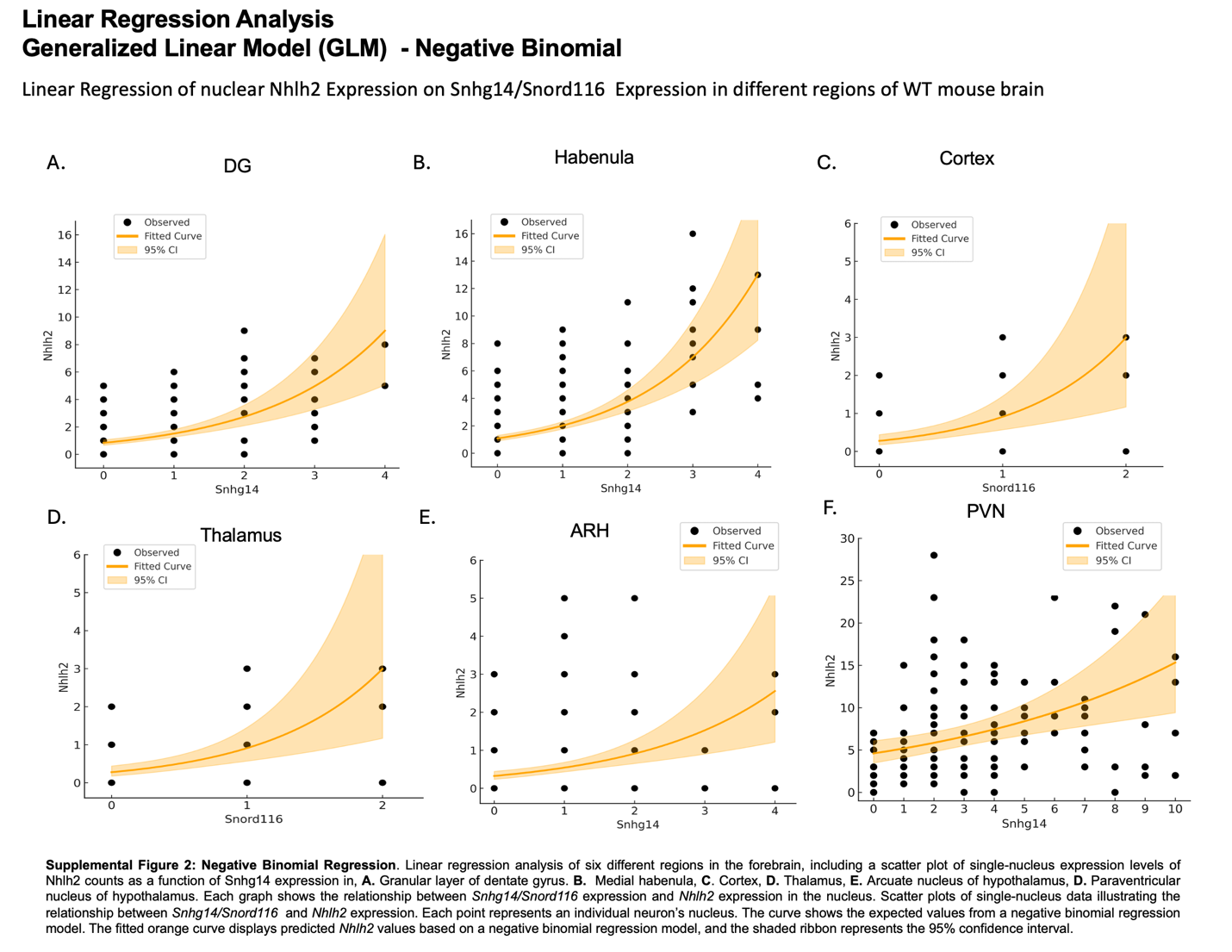
