## Supplemental Video for "Subcellular partitioning of *Nhlh2* mRNA reveals how *SNORD116* loss contributes to Prader-Willi Syndrome"

*Video File: https://drive.google.com/file/d/127vI96YeSGaS16g07UgrNMg76FMe9TZu/view?*

*usp=sharing*

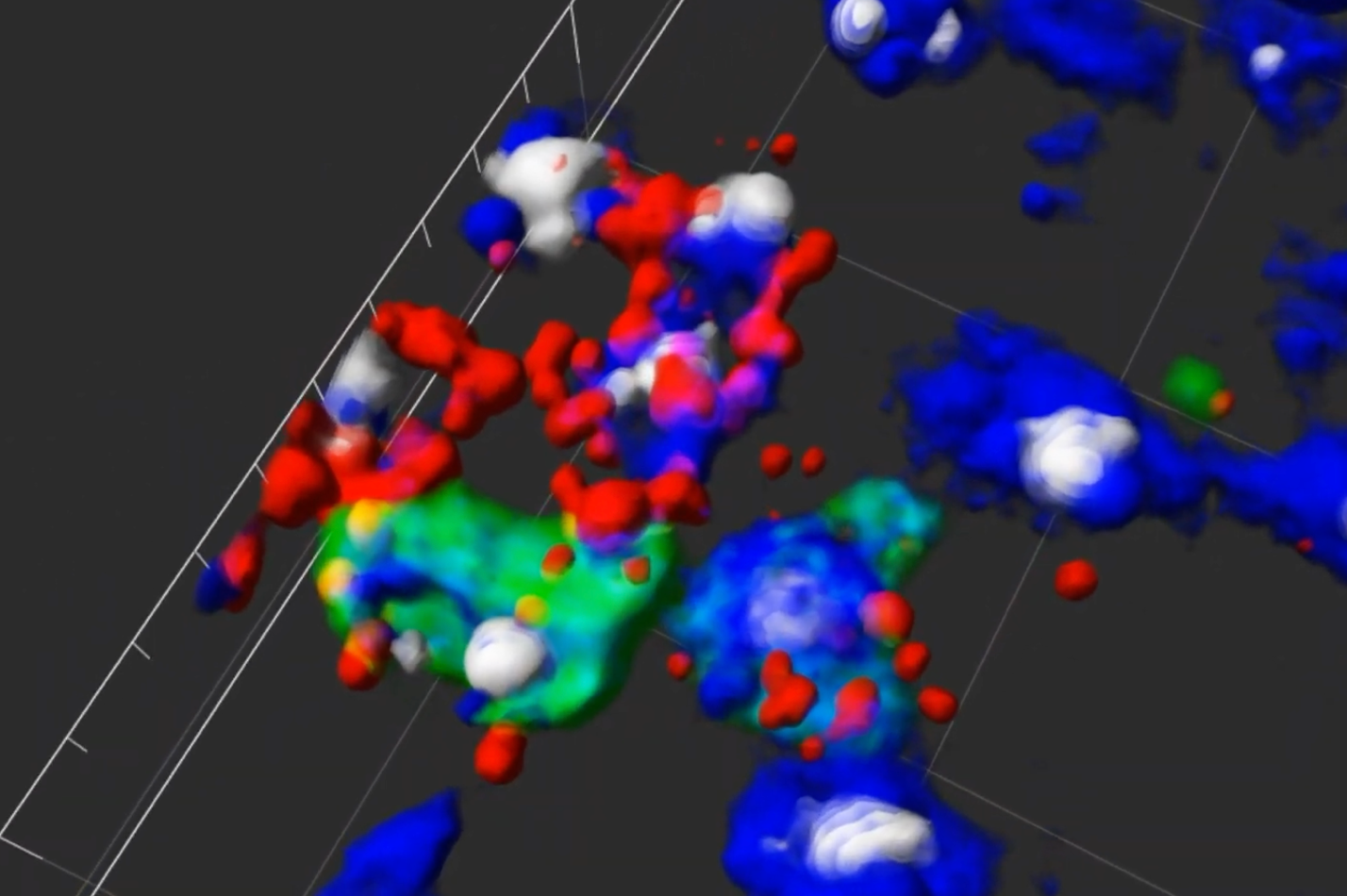
<https://drive.google.com/file/d/127vI96YeSGaS16g07UgrNMg76FMe9TZu/view?%20usp=sharing>
